# smCounter2: an accurate low-frequency variant caller for targeted sequencing data with unique molecular identifiers

**DOI:** 10.1101/281659

**Authors:** Chang Xu, Xiujing Gu, Raghavendra Padmanabhan, Zhong Wu, Quan Peng, John DiCarlo, Yexun Wang

**Affiliations:** Life Science Research and Foundation, QIAGEN Sciences, Inc. 6951 Executive Way, Frederick, Maryland 21703, USA

**Keywords:** low-frequency mutation, unique molecular identifier, DNA polymerase error, targeted sequencing, homopolymer, variant caller

## Abstract

**Motivation:** Low-frequency DNA mutations are often confounded with technical artifacts from sample preparation and sequencing. With unique molecular identifiers (UMIs), most of the sequencing errors can be corrected. However, errors before UMI tagging, such as DNA polymerase errors during end-repair and the first PCR cycle, cannot be corrected with single-strand UMIs and impose fundamental limits to UMI-based variant calling.

**Results:** We developed smCounter2, a UMI-based variant caller for targeted sequencing data and an upgrade from the current version of smCounter. Compared to smCounter, smCounter2 features lower detection limit at 0.5%, better overall accuracy (particularly in non-coding regions), a consistent threshold that can be applied to both deep and shallow sequencing runs, and easier use via a Docker image and code for read pre-processing. We benchmarked smCounter2 against several state-of-the-art UMI-based variant calling methods using multiple datasets and demonstrated smCounter2’s superior performance in detecting somatic variants. At the core of smCounter2 is a statistical test to determine whether the allele frequency of the putative variant is significantly above the background error rate, which was carefully modeled using an independent dataset. The improved accuracy in non-coding regions was mainly achieved using novel repetitive region filters that were specifically designed for UMI data.

**Availability:** The entire pipeline is available at https://github.com/qiaseq/qiaseq-dna under MIT license.

## 1 Introduction

Detection of low-frequency variants is important for early cancer diagnosis and is a very active area of research. Targeted DNA sequencing generates very high coverage over a specific genomic region, therefore allowing low-frequency variants to be observed from a reasonable number of reads. However, distinguishing the observed variants from experimental artifacts is very difficult when the variants’ allele frequencies are near or below the noise level. Providing an error-correction mechanism, unique molecular identifiers (UMIs) have been implemented in several proof-of-concept studies [1–6] and used in translational medical research [7–9]. In these protocols, UMIs (short oligonulceotide sequences) are attached to endogenous DNA fragments by ligation or primer extension, carried along through amplification and sequencing, and finally identified from the reads. Sequencing errors can be corrected by majority vote within a UMI family, because reads sharing a common UMI and random fragmentation site should be identical except for rare collision events [10] or errors within the UMI sequences. DNA polymerase errors occurring during DNA end repair and early PCR cycles (particularly the first cycle), however, cannot be corrected because all reads in the UMI would presumably carry the error. Although PCR error rates are low (10^-4^ - 10^-6^, depending on the enzyme and types of substitution), they impose fundamental limits to UMI-based variant calling.

A two-step UMI-based variant calling approach that first constructs a consensus read with tools like fgbio [11] and then applies one of the conventional low-frequency variant callers [12] to the consensus reads has been implemented in [3, 13]. In addition to the two-stage method, three UMI-based variant callers, DeepSNVMiner [14], smCounter [15], and MAGERI [16], are publicly available. DeepSNVMiner relies on heuristic thresholds to draw consensus and call variants. By default, a UMI is defined as “supermutant” if 40% of its reads support a variant and two supermutants are required to confirm the variant. smCounter was released in 2016 by our group and reported above 90% sensitivity at fewer than 20 false positives per megabase for 1% variants in coding regions. smCounter’s core algorithm consists of a joint probabilistic modeling of PCR and sequencing errors. MAGERI is a collection of tools for UMI-handling, read alignment, and variant calling. The core algorithm estimates the first-cycle PCR errors as a baseline and calls variants whose allele frequencies are higher than the baseline level. MAGERI reported 93% area under curve (AUC) on variants with about 0.1% allele frequencies.

In this article, we present smCounter2, a single nucleotide variant (SNV) and short indel caller for UMI-based targeted sequencing data. smCounter2 offers significant upgrades from its predecessor (smCounter) in terms of algorithm, performance, and usability. smCounter2 adopts the widely popular Beta distribution to model the background error rates and Beta-binomial distribution to model the number of non-reference UMIs. An important feature of smCounter2 is that the model parameters are dynamically adjusted for each input read set. In addition, smCounter2 uses a regression-based filter to reject artifacts in repetitive regions while retaining most of the real variants. The algorithm improvements help to push the detection limit down to 0.5% from the previously reported 1% and increase the sensitivity and specificity compared to other UMI-based methods (two-step consensus-read approach and smCounter), as shown in Section 3. For ease of use, smCounter2 has been released with a Docker container image that includes the complete read processing (using reads from a QIAGEN QIAseq DNA targeted enrichment kit as an example) and variant calling pipeline as well as all the supporting packages and dependencies.

## 2 Methods

### 2.1 smCounter2 workflow

smCounter2’s workflow (Fig. 1) begins with read processing steps that 1) remove the exogenous sequences such as PCR and sequencing adapters and UMI, 2) identify the UMI sequence and append it to the read identifier for downstream analyses, and 3) remove short reads that lack enough endogenous sequence for mapping to the reference genome. The trimmed reads are mapped to the reference genome with BWA-MEM, followed by filtering of poorly mapped reads and soft-clipping of gene-specific primers. A UMI with much smaller read count is combined with a much larger read family if their UMIs are within edit distance of 1 and the corresponding 5’ positions of aligned R2 reads are within 5 bp (i.e. at the random fragmentation site). After UMI clustering, the aligned reads (BAM format) are sent for variant calling. Like many variant callers, smCounter2 walks through the region of interest and processes each position independently. At each position, the covering reads go through several quality filters and the remaining high-quality reads are grouped by putative input molecule (as determined by both the clustered UMI sequence and the random fragmentation site). A consensus base call (including indels) is drawn within a UMI if ≥ 80% of its reads agree. The core variant calling algorithm is built on the estimation of background error rates, i.e. the baseline noise level for the data. A potential variant is identified only if the signal is well above that level (Section 2.2, 2.3). The potential variants are subject to post-filters, including both traditional filters such as strand bias and novel model-based, UMI-specific repetitive region filters (Section 2.4). Finally, the variants are annotated with SnpEff [17] and SnpSift [18] and output in VCF format.

**Figure 1:**
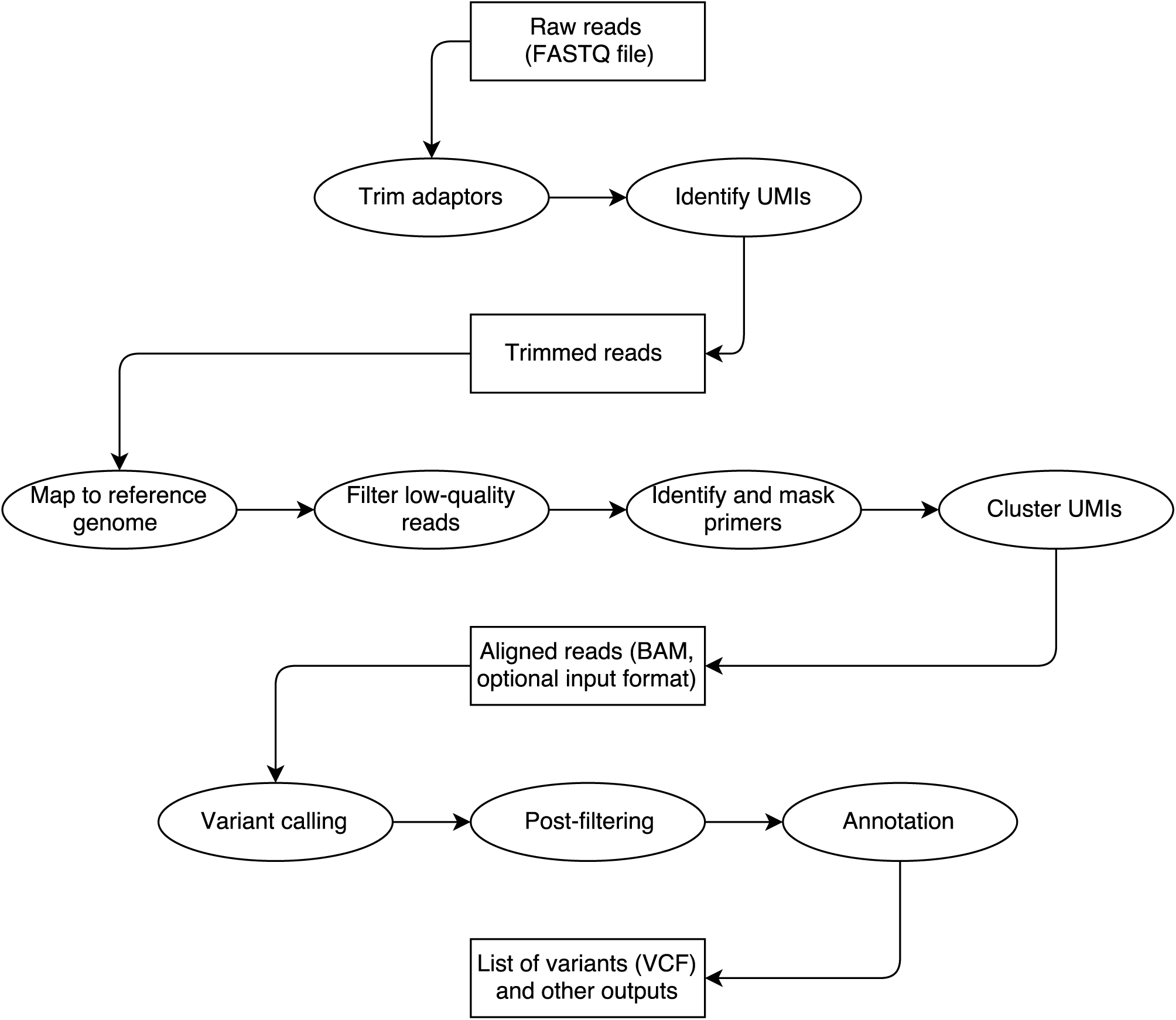
smCounter2 workflow. Rectangular boxes represent the data files and elliptical boxes represent steps of the pipeline. Users can choose to run the whole pipeline from FASTQ to VCF or run the variant calling part only from BAM to VCF.

### 2.2 Estimation of background error rates

Estimating the background error rates is one of the commonly used strategies in somatic variant calling. EBCall [19] and shearwater [20] assume that each site has a distinctive error rate (predominantly sequencing errors) that follows a Beta distribution. LoLoPicker [21] estimates site-specific sequencing error rates as fixed values. For UMI-tagged data, background errors can come from base mis-incorporation by DNA polymerase during end repair and the first-cycle PCR reaction, oxidation damage to DNA bases during sonication shearing and probe hybridization [6, 22], UMI mis-assignment, misalignment, and polymerase slippage (often in repetitive sequences), etc. iDES [6] characterizes the site-specific background error rates in duplex-sequencing data using Normal or Weibull distributions. The limitation of these algorithms is the requirement of many control samples for the site-specific error modeling. As an alternative, MAGERI [16] assumes a universal Beta distribution for all sites, which may result in lower accuracy compared to site-specific error modeling, but as a trade-off requires only one control sample, if the UMI coverage is high enough to observe the background errors and enough sites are covered to reveal the full distribution of error rates.

smCounter2 takes similar experimental and modeling approaches as MAGERI with important modifications. To obtain high-depth data for error profiling, we sequenced 300ng of NA12878 DNA within a 17kbp region using a custom QIAseq DNA panel. After excluding the known SNPs (Genome in a Bottle Consortium [23]), we calculated the error rates by base substitution at each site assuming any non-reference UMIs are background errors. The calculation process is explained in Supplementary Materials, Section 1. We observed notable variation across different base substitutions and that transitions were more error prone than transversions (Fig. 2a). We used the Beta distribution to fit the observed error rates (`R fitdistrplus` [24], Fig. 2b). The quantile plot indicates good fit in general and underestimation of the tail, possibly due to outliers (Fig. 2c). We prepared two versions of error models, one excluding singletons (UMIs with only one read pair) and the other including singletons, to accommodate deep and shallow sequencing depths. For read sets with mean read pair per UMI (rpu) ≥ 3, smCounter2 drops singletons to reduce errors and uses the error model without singletons. For read sets with rpu < 3, smCounter2 keeps some or all singletons (Supplementary Materials, Section 3) to avoid losing too many UMIs and uses the error model with singletons.

**Figure 2:**
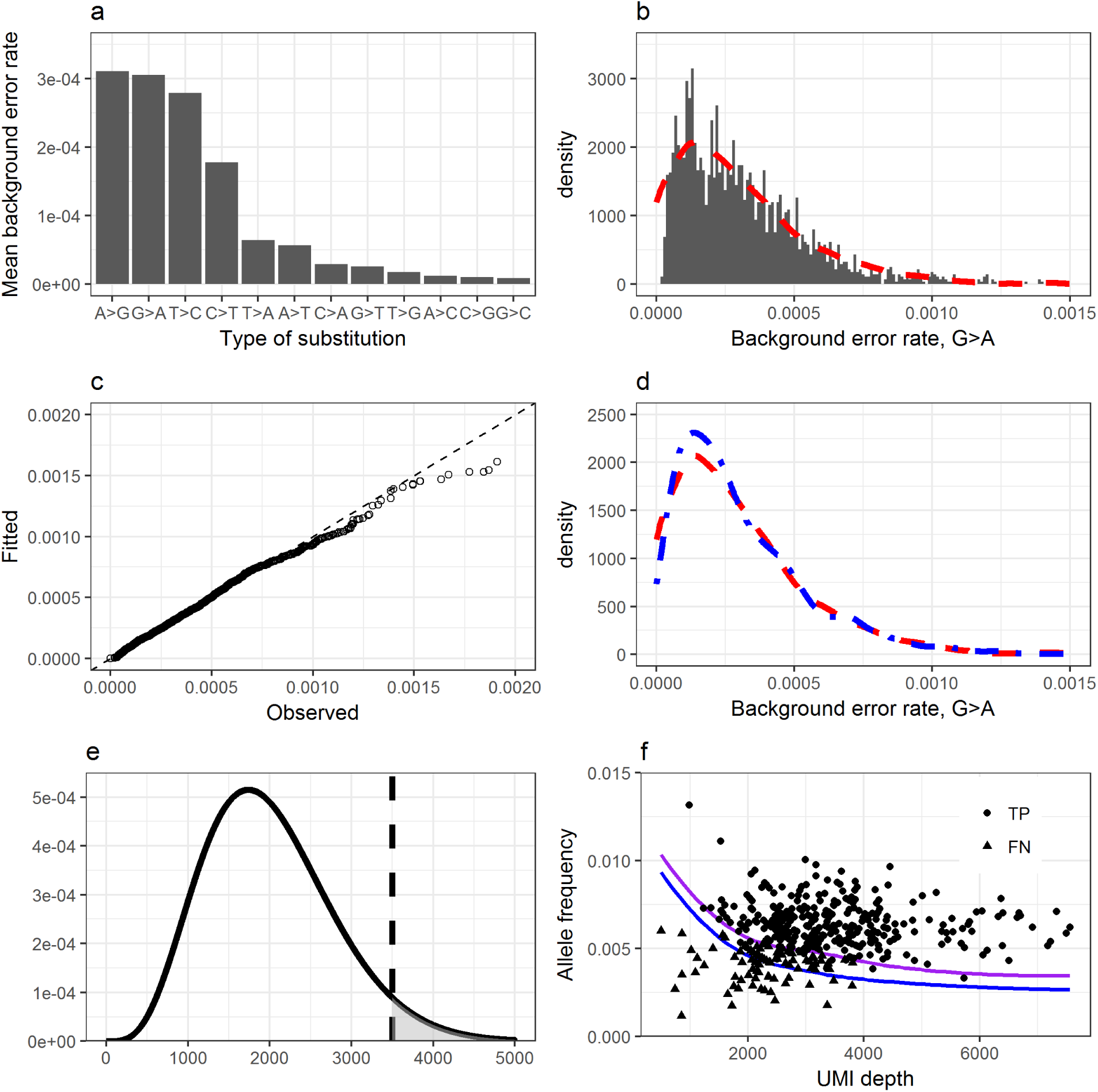
Underlying model of smCounter2. **a** Background error rates for each type of base change, averaged across the panel of M0466. **b** Modeling of the background error rates using the Beta distribution. The histogram shows the frequency of observed G>A error rates in M0466. The red curve is the density of the fitted Beta distribution. **c** Quantile plot to check the goodness-of-fit of the G>A error rate modeling. The observed and fitted quantiles form a 45 degree line in most places, indicating perfect fit. The tail skews towards “observed”, indicating under-estimation of the extremely high error rates. This may simply be explained by outliers, or suggests that a distribution (or mixed distributions) with heavier tail is needed. **d** An real example of parameter adjustment. The red curve is the originally fitted Beta distribution. The blue curve is the adjusted error model with the mean of the input data (N13532) and the original variance. **e** Illustration of the variant calling p-value. The density curve is a hypothesized Beta-binomial distribution. The vertical line indicates the observed non-reference UMI counts. The area of the shaded region is the p-value. **f** Detection limit prediction and confirmation. The purple and blue curves are the predicted site-wise detection limit for Ti and Tv/indels respectively. The dots are the true variants in N13532 (outliers with extremely low UMI depth or high allele frequency excluded). For the dots, the y-axis represents the observed allele frequencies. Round dots are the variants detected and triangle dots are the ones not detected, concentrated in the low enrichment regions.

As a distinctive feature of smCounter2, the Beta distribution parameters are adjusted for each dataset to account for the run-to-run variation. Because the true variants are unknown in the application dataset, we conservatively assumed that all non-reference alleles with VAF below 0.01 are background errors. The low DNA input in most applications impose another challenge in that few of the applications generate enough site-wise UMI coverage for any meaningful update of the error rate distribution. Fortunately, sufficient UMIs can usually be obtained by aggregating the target sites to accurately estimate the mean. Therefore, we only adjust the mean of the Beta distribution to equate the panel-wise mean and leave the dispersion unchanged (Fig. 2d). In specific, the adjusted Beta parameters are

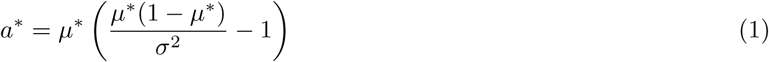

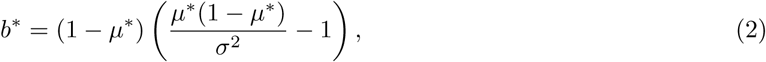

where *µ*\* is the mean error rate of the current data and *σ*^2^ is the variance of the error rate from our control sample. The adjusted distribution *Beta*(*a*\*, *b*\*) has a mean of *µ*\* and variance of *σ*^2^.

Background errors are sensitive to enrichment chemistry and DNA polymerase. The error pattern we observed in QIAseq DNA panels agrees with that in other PCR enrichment studies [25, 26] but differs from hybridization capture studies [6, 22] where A>C and *G* > *T* errors are dominant. Also, certain high-fidelity DNA polymerases have been shown to generate tens- or hundreds-fold lower error rates [25]. Therefore, we did not attempt to build a universal error model by pooling data from multiple experiments with different polymerases as MAGERI did, but instead suggest users who run hybridization capture protocols or use non-QIAseq enrichment chemistry to build their own error profile (a script is provided in the Github repository). Limited by sequencing resources, we were unable to obtain adequate site-wise UMI depth to model base substitutions with low error rates, including all transversions and some transitions. This deficit had several impacts on our modeling procedure. First, we had to assume that all transitions followed the distribution of G>A (second highest) and all transversions followed the distribution of C>T (higher than all transversions). This conservative configuration ensured that the error rates were not under-estimated, but also prevented us from reaching the theoretical detection limit. Second, we were unable to model the indel error rates because 1) indel polymerase errors occur more frequently in repetitive regions, and our panel did not include enough such regions, 2) there are countless types of indels and we cannot model the errors by each type, and 3) indel polymerase error rates are on average lower than base substitution and we lacked the UMI depth to observe enough of them. Again, we conservatively assumed that indel error rates followed the distribution of G>A. Third, because the error rates are very low, zero non-reference UMIs were observed at some sites, especially in low enrichment regions. Depending on the percentage of such sites, we either imputed the zeros with small values or used a zero-inflated Beta distribution (a mixture of Beta distribution and a spike of zeros) instead of Beta.

### 2.3 Statistical model for variant calling and detection limit prediction

We treated variant calling as a hypothesis testing problem, where the null hypothesis (*H*_0_) is that all non-reference UMIs are from background errors and the alternative hypothesis (*H*_*a*_) is that the non-reference UMIs are from the real variant. We assume that there are *n* UMIs covering a site and *k* of them have the same non-reference allele. Under *H*_0_, *k* follows a Binomial distribution *Bin*(*n, p*) where *p* is the background error rate. If *p* follows the Beta distribution with the adjusted parameter *Beta*(*a*\*, *b*\*), the marginal distribution of *k* given *n, a*\*, *b*\* is Beta-binomial. If a zero-inflated Beta distribution is used, *k* has a non-standard marginal distribution. To compute the p-value, we first simulated random samples of *{p*_*i*_, *i* = 1, *…*, *I}* according to the distribution being used. Then for each *p*_*i*_ we computed *P*_*Bin*_(*K* ≥ *k|n, p*_*i*_) based on the Binomial distribution. The p-value represents the probability of observing ≥ *k* non-reference UMIs at a wild-type site (Fig. 2e) and is approximated by

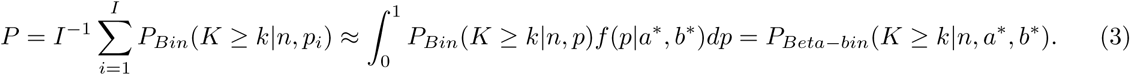

To avoid extremely small fractions, smCounter2 reports *Q* = min(200, - log_10_ *P*) as the variant quality score. The choice of variant calling threshold depends on the tolerance of false positive rate because if the model fits perfectly, the specificity would equal to 1 minus the p-value threshold. By default, smCounter2 aims for ≤ 1 false positives per megabase, which is equivalent to a threshold of *P* ≤ 10^-6^ or *Q* ≥ 6. We will show in Section 3 and Supplementary Materials that this threshold works well for datasets with deep and shallow UMI coverage and for variants with a range of VAFs (0.5, 1, 5% and germlines). The only exception is that, if 0.5-1% indels are of interest, we recommend lowering the Q-threshold to 2.5 to account for the overestimation of indel error rates.

Under this framework, the site-specific detection limit (sDL, the minimum allele frequency to exceed the P-value threshold) is a decreasing function of the UMI depth. It also depends on the type of variant because transitions have higher background error rates than transversions and indels. We estimate that the sDL of transitions is higher than transversions and indels on by about 0.001, or 0.1% in allele frequency. We denote *P* (*n, k, t*) as the p-value given UMI depth *n*, non-reference UMI count *k*, and the type of variant ∈ *{*Ti, Tv+indel*}*. *P* (*n, k, t*) can be computed by Equation (3). The sDL is denoted as arg min_*k*_*{P* (*n, k, t*) < threshold*}/n* and can be computed numerically. Importantly, the predicted sDL is the *observed* allele frequency that often deviates from the true allele frequency in the sample due to random enrichment bias. If we loosely define the overall detection limit as the minimum *true* allele frequency that the variant caller can detect with good sensitivity and specificity, the overall detection limit is usually higher than sDL. Based on our calculation, the theoretical detection limit of a QIAseq DNA panel is around 0.5% when UMI depth is between 2,000 and 4,000. This detection limit was confirmed experimentally by sequencing a sample with known 0.5% variants (Fig. 2f).

### 2.4 Repetitive region filters based on UMI efficiency

Repetitive regions such as homopolymers and microsatellites are enriched in non-coding regions where variants can have important functions from regulating gene expression to promoting diseases [27]. Unfortunately, these regions are a major source of false variant calls due to increased polymerase and mapping errors. For instance, polymerase slippage (one or more bases of the template are skipped over during base extension) occurs more frequently at homopolymers and results in false deletion calls. Reads may be incorrectly mapped to similar regions or mis-aligned if they do not span the whole repetitive sequence, both causing false variant calls. Conventional variant callers apply heuristic filters to remove false calls. For example, Strelka [28] rejects somatic indels at homopolymers with ≥ 8nt or di-nucleotide repeats with ≥ 16nt. Recent haplotype-based variant callers such as GATK HaplotypeCaller [29] perform local *de novo* assembly to avoid mapping/alignment errors in repetitive regions. However, these methods were developed for non-UMI data. smCounter2 includes a set of repetitive region filters that are specifically designed for UMI data. The filters were inspired by the observations that 1) UMIs of the false variants tend to have lower read counts and more heterogeneous reads compared to UMIs of real variants, and 2) reads of the false variants are more likely to contradict with their UMIs’ consensus allele (usually wild-type), whereas reads of the real variants are likely to agree with their UMIs. We used the term “UMI efficiency” to describe these distinctions (Fig. 3a) and quantified the UMI efficiency with four variables: 1) *vafToVmfRatio*, the ratio of allele frequencies based on reads and UMIs; 2) *umiEff*, the proportion of reads that are concordant with their respective UMI consensus; 3) *rpuDiff*, difference of read counts between variant UMIs and wild-type UMIs, adjusted by the standard deviations; and 4) *varRpu*, mean read fragments per variant UMI.

**Figure 3:**
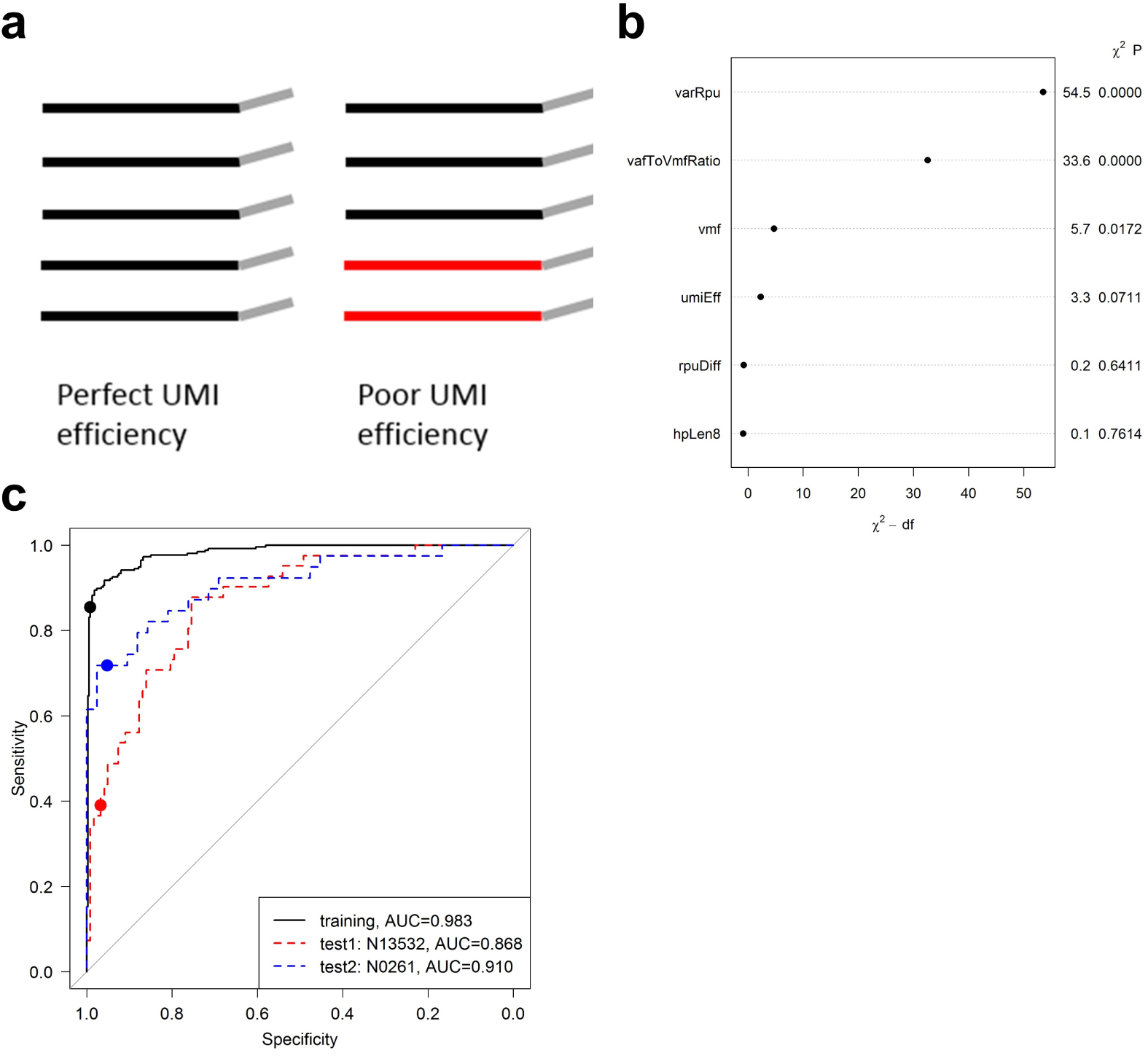
Training and testing of the homopolymer indel filter. **a** Illustration of UMI efficiency. The UMI on the left has perfect efficiency because all reads contributed to the consensus. The UMI on the right has low efficiency because two reads in red disagree with the majority and thus are wasted. smCounter2 requires 80% agreement to reach a consensus, so the entire UMI would be dropped and the other three reads would be wasted as well. **b** Relative importance of each predictor ranked by the explained variation minus the degree of freedom. The read pairs per variant UMI (*varRpu*) and the ratio between allele frequencies by read and by UMI (*vafToVmfRatio*) are the two variables with the most predictive power. The plot is generated with R `rms` package. **c** ROC curves of the logistic regression classifier. The black curve is for the training data that combined all true and false homopolymer indels in N0030, N0015, N11582, and N0164. The blue and red curves are for two test datasets N13532 and N0261, respectively. The dots represent the actual sensitivity and specificity at the cutoff, which is consistent in all three datasets.

We trained and validated a logistic regression model to distinguish real homopolymers indels from artifacts. We focused our resources on this repetitive region subtype because during development, we observed that homopolymer indels were the main contributor of false positives. We combined data from several UMI-based sequencing experiments to assemble a training set with 255 GIAB high confidence homopolymer indels with allele frequencies from 1 to 100% and 386 false positives that would otherwise be called without the filters. In addition to the UMI efficiency variables, we included *sVMF* (VAF based on UMI) and *hpLen8* (binary variable indicating whether the repeat length ≥ 8) as predictors. We found that *varRpb* and *vafToVmfRatio* were the two most important predictors in terms of explained log-likelihood (Fig. 3b). We chose the cutoff on the linear predictors to target on the highest sensitivity while maintaining 99% specificity using the R package OptimalCutpoints [30]. The model and cutoff were applied to two independent datasets N13532 and N0261, both containing 0.5% variants. N13532 had 41 real homopolymer indels and 122 false positives with *Q* ≥ 2.5. The predictive model achieved 39.0% sensitivity, 96.7% specificity, and 0.868 area under the curve (AUC). N0261 had 39 real homopolymer indels and 42 false positives with *Q* ≥ 2.5. The predictive model achieved 71.8% sensitivity, 95.2% specificity, and 0.910 AUC (Fig. 3c).

For other subtypes of variants and repetitive regions, we used heuristic thresholds as filters due to lack of training data. The model parameters and default thresholds are presented in the Supplementary Materials.

## 3 Results

### 3.1 Training and validation datasets

To develop the statistical model and fine-tune the parameters, we did multiple sequencing runs using reference materials NA12878 and NA24385, both of which have high-confidence variants released by GIAB (v3.3.2 used for this study). We mixed small amounts of NA12878 DNA into NA24385 based on the amount of amplifiable DNA measured by QIAseq DNA QuantiMIZE assay to simulate low-frequency variants. The modeling of background error rates was based on M0466, a high-input, deep-sequencing run that reached over 45,000 UMI coverage per site. The selection of variant calling threshold and refinement of filter parameters were based on N0030, N0015, and N11582. N0030 was generated by sequencing 2% mixture of NA12878 on a QIAseq custom DNA panel of 194 cancer-related genes (catalog number CDHS-13026Z-10867). N0015 was based on 10% mixture of NA12878 on a custom panel focusing on non-coding regions (catalog number CDHS-13244Z-3587). Both datasets were also used for the development of smCounter and described in [15]. N11582 was generated with pure NA24385 using QIAseq Human Inherited Disease Panel (catalog number CDHS-14433Z-11582). The main purpose of this dataset was to confirm smCounter2’s ability to call germline variants. Finally, to develop the repetitive region filters, we generated N0164 by pooling reads from 6 libraries with 1, 2.5, 5, 10, 15, and 20% NA12878 admixtures on a custom panel that targeted homopolymer indels.

After development, we tested smCounter2 on independent datasets without any modification to the algorithm and parameters. The first test data N13532 was generated with 1% mixture of NA12878 because the predicted detection limit of our assay’s detection was around 0.5% (Section 2.3). To minimize bias, we designed a new 228-gene custom panel that did not overlap with any of the training datasets (catalog number CDHS-13532Z-10181). Because N13532 did not have enough heterozygous NA12878 indels (164 in the panel, 49 in coding regions), we designed a new panel to capture 269 indels (naive from the training sets and N13532, catalog number CDHS-13907Z-562) and sequenced the 1% NA12878 admixture using that panel to generate the second test data N0261. Strictly speaking, both datasets were not completely independent from the training data because the same DNA samples were used. Therefore, we prepared the third test data M0253 by mixing Horizon Dx’s Tru-Q 7 reference standard (verified 1.3%-tier variants) with Tru-Q 0 (wild-type) at 1:1 ratio. The Horizon sample was sequenced using QIAseq Human Actionable Solid Tumor Panel (catalog number DHS-101Z).

The datasets involved in this study are summarized in Table 1.

**Table 1:** Key statistics of the datasets used for training and testing of smCounter2.

| Dataset | Purpose | Sample | Target re-<br>gion (bp) | Mean<br>UMI<br>depth | Mean<br>read pairs<br>per UMI | VAF (%) | SNVs | indels |
| --- | --- | --- | --- | --- | --- | --- | --- | --- |
| M0466 | training | 0.2%<br>NA12878 | 17,859 | 45,335 | 3.2 | 0.1 | 87 | 0 |
| N0030 | training | 2%<br>NA12878 | 1,032,301 | 3,612 | 8.6 | 1 | 363 | 56 |
| N0015 | training | 10%<br>NA12878 | 406,846 | 4,825 | 8.5 | 5 | 4,412 | 369 |
| N11582 | training | 100%<br>NA24385 | 1,094,204 | 479 | 2.6 | 50 or 100 | 729 | 49 |
| N0164 | training | 1-20%<br>NA12878 | 66,661 | 3,692 | 11.5 | 0.5-10 | 237 | 177 |
| N13532 | test | 1%<br>NA12878 | 928,315 | 4,040 | 7.6 | 0.5 | 293 | 164 |
| N0261 | test | 1%<br>NA12878 | 45,299 | 3,384 | 13.8 | 0.5 | 5 | 269 |
| M0253 | test | 50% HDx<br>Tru-Q 7 | 38,370 | 4,980 | 13.0 | $\geq 0.5$ | 36 (with<br>MNPs) | 1 |

### 3.2 Benchmarking 0.5% variant callling performance using mixed GIAB samples

We benchmarked smCounter2 against four state-of-the-art UMI variant calling algorithms (fgbio+MuTect, fgbio+VarDict, MAGERI, and smCounter) on N13532, which contained 0.5% NA12878 variants. The first two algorithms represent the two-step approach discussed in Section 1. We first constructed consensus reads from the aligned reads (BAM file) using fgbio’s `CallMolecularConsensusReads` and `FilterConsensusReads` functions and then applied two popular low-frequency variant callers, MuTect [31] and VarDict [32], on the consensus reads. MAGERI and smCounter are two representative UMI-aware variant callers. The results (Fig. 4), stratified by type of variant (SNV and indel) and genomic region (all, coding, and non-coding), were measured by sensitivity and false positives per megabase (FP/Mbp, or 10^6^(1 - specificity)) at several thresholds. smCounter2 outperformed the other methods in all categories. In coding regions, smCounter2 achieved 92.4% sensitivity at 12 FP/Mbp for SNVs and 84.4% sensitivity at 7 FP/Mbp for indels (Table 2). In non-coding regions, smCounter2 was able to maintain comparable accuracy for SNVs (83.3% sensitivity at 4 FP/Mbp), but produced lower sensitivity (56.8%) and higher false positive rate (42 FP/Mbp) for indels. In the indel-enriched dataset N0261, smCounter2 produced consistent sensitivity (81.4% in coding and 61.3% in non-coding) and seemingly higher FP/Mbp (0 in coding and 114 in non-coding). However, FP/Mbp in N0261 was based on a very small target region (45kbp) and therefore provides a less accurate specificity estimate.

**Table 2:** smCounter2 performance in detecting 0.5, 1, 5, and 50-100% variants, stratified by type of variant (SNV and indel) and genomic region (coding and non-coding). The metrics were generated with the default thresholds (*Q* ≥ 2.5 for indels in N13532, N0261, M0253 and *Q* ≥ 6 for all other cases). The allele frequency and the purpose of the dataset are displayed under the dataset name. All performance metrics are measured on GIAB high-confidence regions only, the sizes of which are presented in the last column.

| Dataset | Region | Type | TP | FP | FN | TPR(%) | FP/Mbp | PPV(%) | HC size(bp) |
| --- | --- | --- | --- | --- | --- | --- | --- | --- | --- |
| N13532<br>(0.5%, test) | coding | SNV | 171 | 7 | 14 | 92.4 | 12 | 96.1 | 591,154 |
|  |  | indel | 38 | 4 | 7 | 84.4 | 7 | 90.5 | 591,154 |
|  | non-coding | SNV | 90 | 1 | 18 | 83.3 | 4 | 98.9 | 259,162 |
|  |  | indel | 67 | 11 | 51 | 56.8 | 42 | 85.9 | 259,162 |
| N0261<br>(0.5%, test) | coding | indel | 35 | 0 | 8 | 81.4 | 0 | 100.0 | 6,119 |
|  | non-coding | indel | 138 | 4 | 87 | 61.3 | 114 | 97.2 | 35,172 |
| M0253<br>(0.5-30%, test) | all | SNV/MNV | 32 | - | 4 | 88.9 | - | - | 38,370 |
|  |  | indel | 0 | - | 1 | 0.0 | - | - | 38,370 |
| N0030<br>(1%, training) | coding | SNV | 214 | 5 | 4 | 98.2 | 7 | 97.7 | 694,189 |
|  |  | indel | 36 | 1 | 3 | 92.3 | 1 | 97.3 | 694,189 |
|  | non-coding | SNV | 137 | 3 | 8 | 94.5 | 13 | 97.9 | 236,687 |
|  |  | indel | 12 | 3 | 5 | 70.6 | 13 | 80.0 | 236,687 |
| N0015<br>(5%, training) | coding | SNV | 528 | 0 | 4 | 99.2 | 0 | 100.0 | 35,718 |
|  |  | indel | 9 | 0 | 1 | 90.0 | 0 | 100.0 | 35,718 |
|  | non-coding | SNV | 3851 | 7 | 29 | 99.3 | 24 | 99.8 | 297,805 |
|  |  | indel | 285 | 13 | 74 | 79.4 | 44 | 95.6 | 297,805 |
| N11582<br>(50-100%,<br>training) | coding | SNV | 421 | 2 | 0 | 100.0 | 3 | 99.5 | 682,483 |
|  |  | indel | 4 | 0 | 0 | 100.0 | 0 | 100.0 | 682,483 |
|  | non-coding | SNV | 301 | 1 | 7 | 97.7 | 4 | 99.7 | 269,761 |
|  |  | indel | 34 | 1 | 11 | 75.6 | 4 | 97.1 | 269,761 |

**Figure 4:**
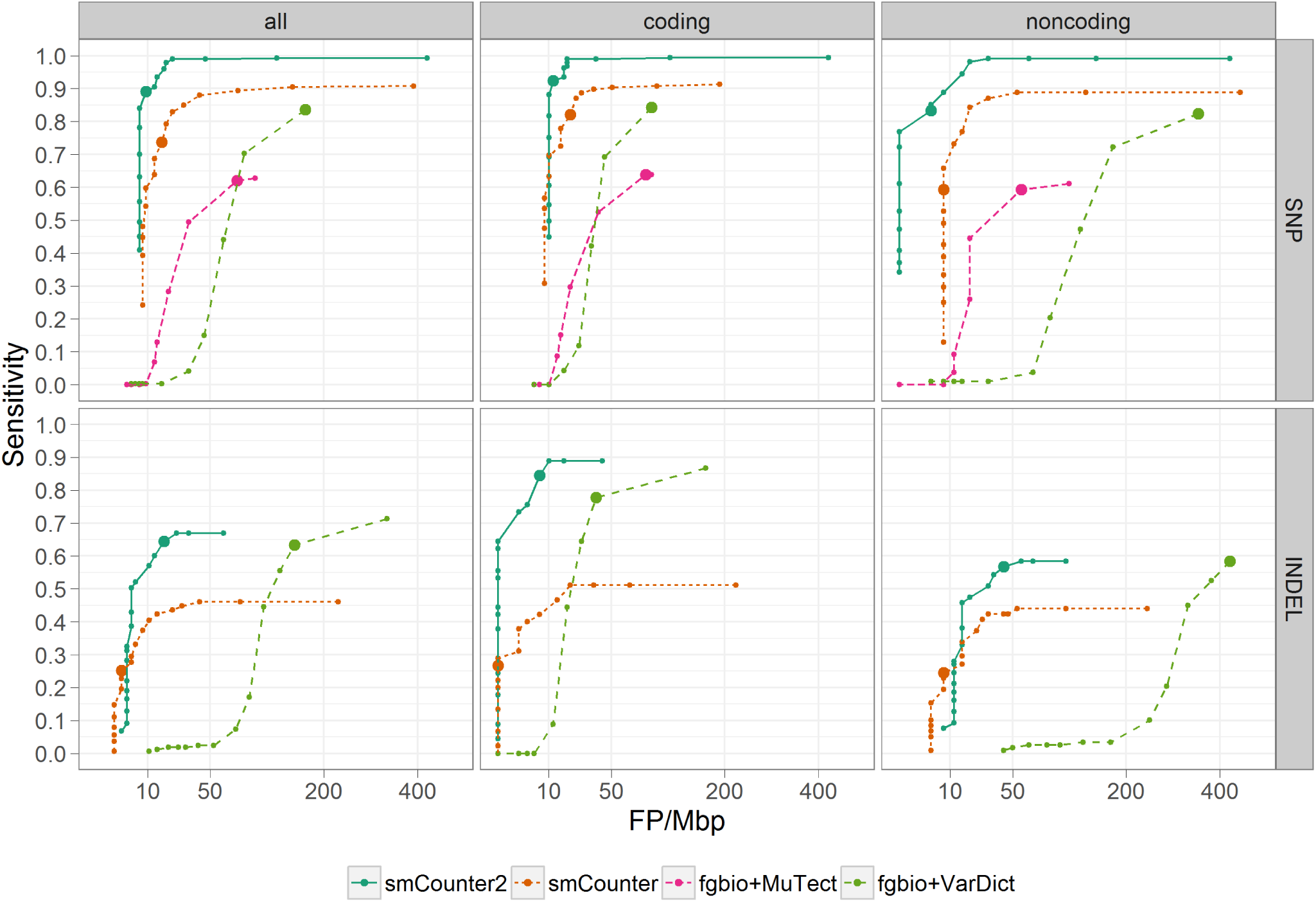
Benchmarking smCounter2, smCounter, fgbio+MuTect, fgbio+VarDict on 0.5% variants in N13532. The performance is measured by false positives per megabase (x-axis) and sensitivity (y-axis), stratified by type of variant (SNV and indel) and region (coding, non-coding, and all). The ROC curves are generated by varying the threshold for each method: Q-score for smCounter2, prediction index for sm-Counter, likelihood ratio for MuTect, and minimum allele frequency for VarDict. MuTect does not detect indels so is not included in the indel comparison.

We did not show MAGERI’s performance in Fig. 4 because it is unfair to compare MAGERI with smCounter2 using QIAseq data. MAGERI’s error model is based only on primer extension assays from a mix of DNA polymerases including several high-fidelity enzymes [26], while smCounter2’s error model is specific to the entire QIAseq targeted DNA panel workflow, including DNA fragmentation, end repair and PCR enrichment steps. Because the MAGERI error model does not include errors introduced at the typical DNA fragmentation and end repair process (their assays do not have those steps), MAGERI’s background error rates are lower than those in smCounter2. For example, the mean error rate of A>G and T>C used by MAGERI is 6.3 × 10^-5^ per base (https://github.com/mikessh/mageri-paper/blob/master/error_model/basic_error_model.pdf) and about 3 × 10^-4^ per base for smCounter2 (Fig. 2a). Therefore, with QIAseq data, MAGERI will produce more false positives due to underestimation of the error rate. We included MAGERI’s ROC curve in Fig. S2 to illustrate the point that the error models are specific to each NGS workflow and need to be empirically established for different workflows.

We used the default setting for smCounter and adjusted the parameters of fgbio, MuTect, and VarDict based on our experience of working with them. However, given the infinite parameter space, we cannot claim that the results reported here reflect their optimal performance. Several variant calling thresholds were used to investigate the sensitivity-specificity trade-off and draw the ROC curves. For fgbio+MuTect, we used MuTect’s likelihood ratio score (LOD score) as threshold. For fgbio+VarDict, we set VarDict’s minimum allele frequency (-f). For MAGERI, we did not use the seemingly obvious threshold “Q-score” because they were not allowed to exceed 100 for computational reasons, and even a Q-score of 100 was overly sensitive and generated too many false calls. Instead, we held Q-score constant at 100 and varied the number of reads in a UMI (-defaultOverseq). The parameters and thresholds used in this study are listed in the Supplementary Materials, Section 3.

### 3.3 Detecting ≥ 1% variants in (possibly) shallow sequencing runs

smCounter2 achieved good sensitivity on 1, 5, 50, and 100% variants as well (Table 2, datasets N0030, N0015, N11582). The biggest advantage for smCounter2 was in non-coding regions due to the repetitive region filters. Compared to smCounter, for 1% non-coding variants, smCounter2’s sensitivity increased from 75.2 to 94.5% for SNVs and from 23.5 to 70.6% for indels (Fig. S3). For 5% non-coding variants, smCounter2’s sensitivity increased from 95.1 to 99.3% for SNVs and from 58.2 to 79.4% for indels (Fig. S4). For 50 and 100% non-coding variants, smCounter2’s sensitivity increased from 89.0 to 97.7% for SNVs and from 42.2 to 75.6% for indels (Fig. S5). Both smCounter2 and smCounter outperformed fgbio+MuTect and fgbio+VarDict on 1 and 5% variants in all categories. For germline variants, however, smCounter2 had lower sensitivity for non-coding indels compared to fgbio+HaplotypeCaller (75.6% vs. 88.9%). This demonstrated the advantage of a haplotype-based strategy in difficult regions. Other than for non-coding indels, the two methods achieved comparable accuracy in other categories.

To test smCounter2’s robustness under low sequencing capacity, we *in silico* downsampled N0030 to 80, 60, 40, 20, and 10% of reads to mimic a range of sequencing and UMI depths. smCounter2 outperformed other methods in all sub-samples (Fig. S6-S10). The downsample series also demonstrated that smCounter2’s constant threshold can maintain consistently low false positive rates for SNV across a range of UMI depths (Fig. 5). In contrast, smCounter’s default threshold must move linearly with the UMI depth to maintain a certain level of false positive rate. Similarly, MuTect’s threshold based on the likelihood ratio needs to be adjusted for datasets with varying read depth. smCounter2’s invariant threshold allows users to apply the default setting to a wide range of sequencing and sample input conditions.

**Figure 5:**
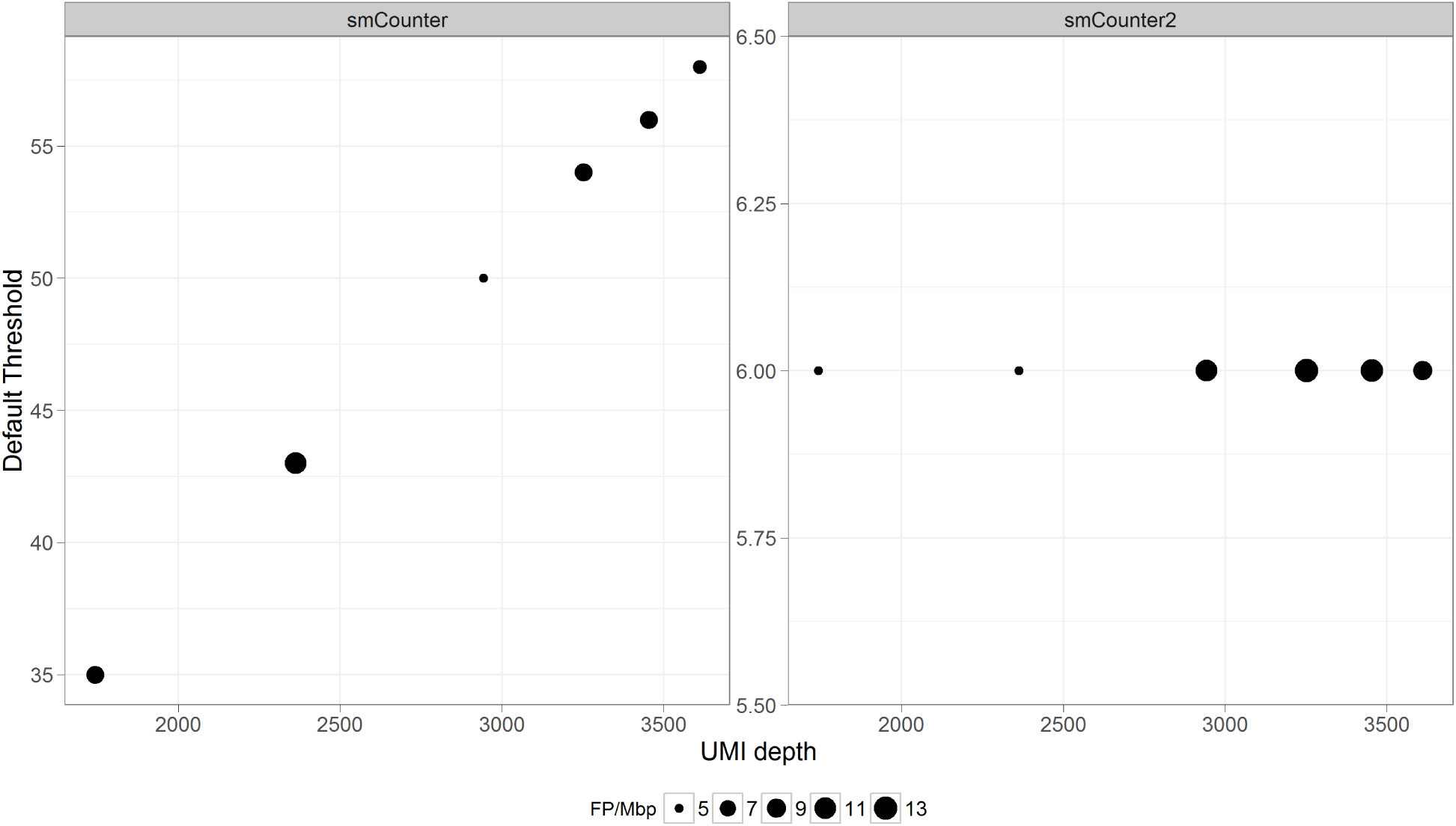
Default thresholds of smCounter and smCounter2 at different UMI depths and associated false positive rates, based on the downsample series of N0030. smCounter’s threshold moves linearly with the UMI depth and is determined using an empirical formula *y* = 14 + 0.012*x*. smCounter2’s threshod is constant at 6. The false positive rates for SNV are well controlled (between 5 to 13 FP/Mbp, represented by the point size) using both methods.

It is important to note that the results described in Section 3.2 and 3.3 are measured over GIAB high confidence region. smCounter2’s performance in GIAB-difficult regions is unknown, both absolutely and in comparison to other variant callers. We also note that the results in Section 3.3 are based on training datasets only. We have not tested smCounter2 on independent 1% or above variants.

### 3.4 Detecting complex cancer mutations using Horizon Tru-Q samples

The performance data described so far were based on diluted NA12878 or pure NA24385, all of which contained germline variants. To test smCounter2 on low-frequency cancer mutations, we sequenced the Tru-Q 7 reference standard (Horizon Dx) that contained verified 1.0% (and above) onco-specific mutations. The sample was diluted 1:1 in Tru-Q 0 (wildtype, Horizon Dx) to simulate 0.5% variants. For this dataset (M0253), smCounter2 detected 32 out of 36 SNV/MNVs (88.9%) and narrowly missed the only deletion (Q=2.49 for threshold of 2.5). Because not all variants in the Tru-Q samples are known, we cannot evaluate specificity using this dataset. The list of variants in this dataset, along with the observed VAF and smCounter2 results, can be found in Supplementary File `Ground_truth_variants.xlsx`.

The Tru-Q sample contains some complex multi-allelic variants that are challenging for variant callers that are not haplotype-aware. For example, there are four variants A>C, A>T, AC>CT, and AC>TT at one position (chr7:140453136, GRCh37) and a C>T point mutation at the next position. smCounter2 detected the three SNVs but failed to recognize the two MNVs.

## 4 Discussion

In this paper, we described smCounter2, the next version of our UMI-based variant caller. Compared to the previous version of smCounter, smCounter2 features lower detection limit, higher accuracy, consistent threshold, and better usability. smCounter2 pushed the detection limit of QIAseq targeted DNA panels down to 0.5% and achieved over 92% sensitivity for SNVs and 84% for indels in coding regions, at the cost of about 10 false positives per megabase. This result was significantly better than smCounter and other state-of-the-art UMI-based variant callers we benchmarked. smCounter2 achieved a lower detection limit because the background error rates were accurately estimated for specific base incorporation errors. The statistical model allows smCounter2 to quantify the deviation from real variants to the background errors using p-values. Therefore, the ambiguous variants whose allele frequencies are close to the background error rates can be called by smCounter2 with reasonable confidence. Importantly, 0.5% is a not an algorithm limit, but rather a chemistry limit. We believe that smCounter2 can achieve even lower detection limits for other chemistry with lower background error rate.

smCounter2 has higher accuracy than its predecessor for both SNVs and indels, in both coding and non-coding regions, for both deep and shallow sequencing runs, and for both low-frequency (≥ 0.5%) and germline variants. The accuracy improvement is due to the modeling of background error rates and, particularly in non-coding regions, UMI-based repetitive region filters. The filters catch false positives in the repetitive regions that pass the p-value threshold but have low “UMI efficiency”, a novel concept that we have proved to be useful in distinguishing real variants from artifacts. Particularly for indels in homopolymers, smCounter2 employs a logistic regression classifier that was trained and validated with separate datasets.

smCounter2 has a more consistent variant calling threshold (*Q* ≥ 2.5 for 0.5-1% indels and *Q* ≥ 6 for other cases) that is independent from the UMI depth, unlike smCounter or MuTect whose optimal threshold must move with the UMI or read depth. This is because smCounter evaluates potential variants by the *number* of non-reference UMIs, while smCounter2 evaluates potential variants by the *proportion* of non-reference UMIs. Moreover, because smCounter2 performs a statistical test at each site, UMI depth has already been accounted for in the p-value. A higher UMI depth will result in better power of detection without raising the threshold. The consistent threshold makes it easier to benchmark smCounter2 with independent datasets. As pointed out by [12], benchmarking studies face the challenge of tuning the variant callers for different datasets.

smCounter2 is also easier to use than smCounter. The read-processing code has been released together with the variant caller, making smCounter2 a complete pipeline from FASTQ to VCF. Some users may prefer to use their own read-processing script because read structures may differ from protocol to protocol. These users can run the variant caller only with the BAM file as input, if UMIs are properly tagged in the BAM. smCounter2 is released as a Docker container image so that users do not need to install the dependencies manually.

smCounter2 has advantages over other UMI-based variant callers as well. Compared to the two-stage approach, smCounter2 requires less tuning and achieves better detection accuracy with low-frequency variants. In contrast to MAGERI’s strategy of pooling data from several polymerases, smCounter2’s error model is developed using a single dataset with very deep coverage. Library preparation method and DNA polymerase have a large impact on the background error rates. Therefore we believe that profiling the errors per individual polymerase and protocol is a better approach. Furthermore, smCounter2 adjusts the error model for each individual dataset, making it a Bayesian-like procedure where the final error model is determined by both the prior knowledge and the data.

smCounter2 has several limitations. First, the error model is specific to the QIAseq targeted panel sequencing protocol, which uses integrated DNA fragmentation plus end repair process and single primer PCR enrichment. Without further tests, we are less certain if the error model holds for other types of library preparation and enrichment protocols. We are more certain, however, that our error model would not fit the data generated by hybridization capture enrichment due to distinct base errors from hybridization chemistry. We have released the modeling code and encourage users, who want to use smCounter2 on non-QIAseq panel data, to re-estimate the background error rates if datasets with sufficient UMI depth are available. Second, limited by resources, we were not able to generate data with enough UMI depth to accurately estimate the transversion and indel error rates. This deficit prevented the variant caller from reaching the assay’s theoretical detection limit. However, as we continue to generate data, we will update the error models with more precise parameters. Third, the germline indel calling accuracy, especially in non-coding regions, is lower than the two-step approach of fgbio+HaplotypeCaller. Although smCounter2 has very efficient repetitive region filters, it still adopts a base-by-base variant calling strategy and relies on the mapping, which is error-prone in repetitive regions. Haplotype-aware variant callers such as HaplotypeCaller are more effective in repetitive and variant-dense regions because they perform local assembly and no longer rely on the local reference genome alignment information. Fourth, smCounter2 has difficulty in handling very complex variants. For example, it failed to report all minor alleles of the complex, multi-allelic variant in Section 3.4. This can potentially be solved by including haplotype-aware features. We have not tested smCounter2’s reliability in detecting variants with 3 or more minor alleles, partly because these variants are not observed frequently. By default, smCounter2 reports bi- and tri-allelic variants only. Fifth, the benchmarking study was based on reference standards. We have not demonstrated smCounter2’s performance using real tumor samples and therefore cannot claim clinical utility. We hope smCounter2 will be used in both translational and clinical studies and look forward to feedback from users.

## 5 Additional files and availablity of data

`Ground_truth_variants.xlsx` contains high-confidence heterozygous NA12878-not-NA24385 variants (GIAB v3.3.2) in N13532, N0261, N0030, N0015, high-confidence NA24385 variants in N11582, and verified Tru-Q 7 variants in M0253. `Dataset_download_URLs.xlsx` contains the download URLs for the FASTQ and BAM files on Google Cloud Platform. Code and supporting files for background error modeling and benchmarking against GIAB ground truth set are available at https://github.com/qiaseq/smcounter-v2-paper.

## References

[1] Cassandra B Jabara, Corbin D Jones, Jeffrey Roach, Jeffrey A Anderson, and Ronald Swanstrom. Accurate sampling and deep sequencing of the hiv-1 protease gene using a primer id. Proceedings of the National Academy of Sciences, 108(50):20166–20171, 2011.

[2] Michael W Schmitt, Scott R Kennedy, Jesse J Salk, Edward J Fox, Joseph B Hiatt, and Lawrence A Loeb. Detection of ultra-rare mutations by next-generation sequencing. Proceedings of the National Academy of Sciences, 109(36):14508–14513, 2012.

[3] Quan Peng, Ravi Vijaya Satya, Marcus Lewis, Pranay Randad, and Yexun Wang. Reducing amplification artifacts in high multiplex amplicon sequencing by using molecular barcodes. BMC genomics, 16(1):589, 2015.

[4] Yoji Kukita, Ryo Matoba, Junji Uchida, Takuya Hamakawa, Yuichiro Doki, Fumio Imamura, and Kikuya Kato. High-fidelity target sequencing of individual molecules identified using barcode sequences: de novo detection and absolute quantitation of mutations in plasma cell-free dna from cancer patients. DNA Research, 22(4):269–277, 2015.

[5] Scott R Kennedy, Michael W Schmitt, Edward J Fox, Brendan F Kohrn, Jesse J Salk, Eun Hyun Ahn, Marc J Prindle, Kawai J Kuong, Jiang-Cheng Shen, Rosa-Ana Risques, et al. Detecting ultralow-frequency mutations by duplex sequencing. Nature protocols, 9(11):2586–2606, 2014.

[6] Aaron M Newman, Alexander F Lovejoy, Daniel M Klass, David M Kurtz, Jacob J Chabon, Florian Scherer, Henning Stehr, Chih Long Liu, Scott V Bratman, Carmen Say, et al. Integrated digital error suppression for improved detection of circulating tumor dna. Nature biotechnology, 34(5):547–555, 2016.

[7] Andrew L Young, Grant A Challen, Brenda M Birmann, and Todd E Druley. Clonal haematopoiesis harbouring aml-associated mutations is ubiquitous in healthy adults. Nature communications, 7:12484, 2016.

[8] Daniel Z Bar, Martin F Arlt, Joan F Brazier, Wendy E Norris, Susan E Campbell, Peter Chines, Delphine Larrieu, Stephen P Jackson, Francis S Collins, Thomas W Glover, et al. A novel somatic mutation achieves partial rescue in a child with hutchinson-gilford progeria syndrome. Journal of medical genetics, 54(3):212–216, 2017.

[9] Rocio Acuna-Hidalgo, Hilal Sengul, Marloes Steehouwer, Maartje van de Vorst, Sita H Vermeulen, Lambertus ALM Kiemeney, Joris A Veltman, Christian Gilissen, and Alexander Hoischen. Ultra-sensitive sequencing identifies high prevalence of clonal hematopoiesis-associated mutations throughout adult life. The American Journal of Human Genetics, 101(1):50–64, 2017.

[10] Richard H. Liang, Theresa Mo, Winnie Dong, Guinevere Q. Lee, Luke C. Swenson, Rosemary M. McCloskey, Conan K. Woods, Chanson J. Brumme, Cynthia K.Y. Ho, Janke Schinkel, Jeffrey B. Joy, P. Richard Harrigan, and Art F.Y. Poon. Theoretical and experimental assessment of degenerate primer tagging in ultra-deep applications of next-generation sequencing. Nucleic Acids Research, 42(12):e98, 2014.

[11] Fgbio. https://github.com/fulcrumgenomics/fgbio.

[12] Chang Xu. A review of somatic single nucleotide variant calling algorithms for next-generation sequencing data. Computational and Structural Biotechnology Journal, 16:15–24, 2018.

[13] Brendan Blumenstiel, Mark Fleharty, Matthew Defelice, Lisa Green, Jonna Grimsby, Yossi Farjoun, Niall Lennon, and Stacey Gabriel. Understanding low allele variant detection in heterogeneous samples, required read coverage and the utility of unique molecular indices (umis). 2017.

[14] T Daniel Andrews, Yogesh Jeelall, Dipti Talaulikar, Christopher C Goodnow, and Matthew A Field. Deepsnvminer: a sequence analysis tool to detect emergent, rare mutations in subsets of cell populations. PeerJ, 4:e2074, 2016.

[15] Chang Xu, Mohammad R Nezami Ranjbar, Zhong Wu, John DiCarlo, and Yexun Wang. Detecting very low allele fraction variants using targeted dna sequencing and a novel molecular barcode-aware variant caller. BMC genomics, 18(1):5, 2017.

[16] Mikhail Shugay, Andrew R Zaretsky, Dmitriy A Shagin, Irina A Shagina, Ivan A Volchenkov, Andrew A Shelenkov, Mikhail Y Lebedin, Dmitriy V Bagaev, Sergey Lukyanov, and Dmitriy M Chudakov. Mageri: Computational pipeline for molecular-barcoded targeted resequencing. PLoS computational biology, 13(5):e1005480, 2017.

[17] P. Cingolani, A. Platts, M. Coon, T. Nguyen, L. Wang, S.J. Land, X. Lu, and D.M. Ruden. A program for annotating and predicting the effects of single nucleotide polymorphisms, snpeff: Snps in the genome of drosophila melanogaster strain w1118; iso-2; iso-3. Fly, 6(2):80–92, 2012.

[18] P. Cingolani, V.M. Patel, M. Coon, T. Nguyen, S.J. Land, D.M. Ruden, and X. Lu. Using drosophila melanogaster as a model for genotoxic chemical mutational studies with a new program, snpsift. Frontiers in Genetics, 3, 2012.

[19] Yuichi Shiraishi, Yusuke Sato, Kenichi Chiba, Yusuke Okuno, Yasunobu Nagata, Kenichi Yoshida, Norio Shiba, Yasuhide Hayashi, Haruki Kume, Yukio Homma, et al. An empirical bayesian framework for somatic mutation detection from cancer genome sequencing data. Nucleic acids research, 41(7):e89–e89, 2013.

[20] Moritz Gerstung, Elli Papaemmanuil, and Peter J Campbell. Subclonal variant calling with multiple samples and prior knowledge. Bioinformatics, 30(9):1198–1204, 2014.

[21] Jian Carrot-Zhang and Jacek Majewski. Lolopicker: Detecting low-fraction variants in low-quality cancer samples from whole-exome sequencing data. bioRxiv, page 043612, 2016.

[22] Gahee Park, Joo Kyung Park, Seung-Ho Shin, Hyo-Jeong Jeon, Nayoung KD Kim, Yeon Jeong Kim, Hyun-Tae Shin, Eunjin Lee, Kwang Hyuck Lee, Dae-Soon Son, et al. Characterization of background noise in capture-based targeted sequencing data. Genome biology, 18(1):136, 2017.

[23] Justin M Zook, Brad Chapman, Jason Wang, David Mittelman, Oliver Hofmann, Winston Hide, and Marc Salit. Integrating human sequence data sets provides a resource of benchmark snp and indel genotype calls. Nature biotechnology, 32(3):246, 2014.

[24] Marie Laure Delignette-Muller, Christophe Dutang, et al. fitdistrplus: An r package for fitting distributions. Journal of Statistical Software, 64(4):1–34, 2015.

[25] Vladimir Potapov and Jennifer L Ong. Examining sources of error in pcr by single-molecule sequencing. PloS one, 12(1):e0169774, 2017.

[26] Dmitriy A Shagin, Irina A Shagina, Andrew R Zaretsky, Ekaterina V Barsova, Ilya V Kelmanson, Sergey Lukyanov, Dmitriy M Chudakov, and Mikhail Shugay. A high-throughput assay for quantitative measurement of pcr errors. Scientific Reports, 7(1):2718, 2017.

[27] Ekta Khurana, Yao Fu, Dimple Chakravarty, Francesca Demichelis, Mark A Rubin, and Mark Gerstein. Role of non-coding sequence variants in cancer. Nature Reviews Genetics, 17(2):93, 2016.

[28] Christopher T Saunders, Wendy SW Wong, Sajani Swamy, Jennifer Becq, Lisa J Murray, and R Keira Cheetham. Strelka: accurate somatic small-variant calling from sequenced tumor–normal sample pairs. Bioinformatics, 28(14):1811–1817, 2012.

[29] Mark A DePristo, Eric Banks, Ryan Poplin, Kiran V Garimella, Jared R Maguire, Christopher Hartl, Anthony A Philippakis, Guillermo Del Angel, Manuel A Rivas, Matt Hanna, et al. A framework for variation discovery and genotyping using next-generation dna sequencing data. Nature genetics, 43(5):491, 2011.

[30] Móonica Lóopez-Ratóon, María Xosé Rodríiguez-Álvarez, Carmen Cadarso-Suárez, Francisco Gude-Sampedro, et al. Optimalcutpoints: an r package for selecting optimal cutpoints in diagnostic tests. Journal of statistical software, 61(8):1–36, 2014.

[31] Kristian Cibulskis, Michael S Lawrence, Scott L Carter, Andrey Sivachenko, David Jaffe, Carrie Sougnez, Stacey Gabriel, Matthew Meyerson, Eric S Lander, and Gad Getz. Sensitive detection of somatic point mutations in impure and heterogeneous cancer samples. Nature biotechnology, 31(3):213–219, 2013.

[32] Zhongwu Lai, Aleksandra Markovets, Miika Ahdesmaki, Brad Chapman, Oliver Hofmann, Robert McEwen, Justin Johnson, Brian Dougherty, J Carl Barrett, and Jonathan R Dry. Vardict: a novel and versatile variant caller for next-generation sequencing in cancer research. Nucleic acids research, 44(11):e108–e108, 2016.

