## Supplementary Materials for "smCounter2: an accurate low-frequency variant caller for targeted sequencing data with unique molecular identifiers"

### 1 Counting erroneous UMIs

The first step is to calculate the site-wise error rate, i.e., the proportion of erroneous UMIs at each site. We excluded any site with known variant or outside of the high-confidence region (GIAB, v3.3.2). The remaining sites are presumably wild-type where any non-reference allele is a background error. For simplicity, we assumed that all erroneous UMIs are caused by DNA polymerase error, which is the dominant type of errors. Importantly, polymerase errors may or may not be the same as the base change. For example, an A>G base change could be caused by a T>C polymerase error when the primer binds to the forward strand of the original template. After reverse complement and further amplification, the final product is a double strand DNA with G and C on the forward and reverse strands, appearing as an A>G base change (Fig. S1a). Similarly, an A>G base change could also be caused by an A>G polymerase error when the primer binds to the reverse strand (Fig. S1b). The site-wise polymerase error rate can be calculated by strand. For example, if a site with reference base A is covered by $n_+$ UMIs on the forward strand with $x_+$ G's and $n_-$ UMIS on the reverse strand with $x_-$ G's, the A>G polymerase error rate is $x_-/n_-$ and the T>C polymerase error rate is $x_+/n_+$.

### 2 Repetitive region filters

For putative indels on homopolymers (minimum 6nt repeat) that passed the Q-threshold, a predictive score is calculated as follows:

$$\begin{aligned}\text{Score} = &-1.658 + 0.0474 \times \text{vmf} - 0.868 \times \text{hpLen8} \\ &- 1.530 \times \text{vafToVmfRatio} + 3.018 \times \text{umiEff} - 0.066 \times \text{rpuDiff} + 0.513 \times \text{varRpu}.\end{aligned}$$

All the predictors have been described in the main text Section 2.4. The putative indel will be called if the predictive score is above or equal to 0.565 and rejected otherwise. For all other subtypes of repetitive region variants, including homopolymer SNVs and variants in other difficult regions, smCounter2 applies heuristic filters. If the mean read pairs per UMI (rpu) is at least 4, the putative variant is called if umiEff $> 0.1$ and vafToVmfRatio $< 3.0$ and rpuDiff $< 2.5$. If $1.8 \leq$ rpu $< 4$, the putative variant is called if umiEff $> 0.8$ and vafToVmfRatio $< 2.0$ and rpuDiff $< 1.5$. For runs with rpu $< 1.8$, all repetitive region variants are rejected.

### 3 Parameter settings for smCounter2, smCounter, fgbio, MuTect, VarDict, HaplotypeCaller, and MAGERI

We used the following smCounter2 command for all datasets except M0253, where `maxAltAllele` was set to 3 to allow detection of up to 3 minor alleles.

```
python sm_counter_v2.py --runPath /my/path --bamFile my_aligned_reads.bam --bedTarget
    my_target_region.bed --outPrefix my_output_prefix --nCPU 64 --minBQ 25 --minMQ 50 --hpLen 8 --
    mismatchThr 6.0 --primerDist 2 --mtThreshold 0.8 --rpb mean_read_pairs_per_UMI --primerSide 1
    --minAltUMI 3 --maxAltAllele 2 --refGenome my_reference_genome.fa --srBed SR_LC_SL.full.bed --
    repBed simpleRepeat.full.bed
```

We used the following smCounter command. `mtDrop` was set to 1 to drop the singletons for datasets with high read pairs per UMI, including N13532, N0261, M0253, N0030 and 80, 60, and 40% downsamples, and N0015. `mtDrop` was set to 0 to include the singletons for shallow sequencing runs, including N11582 and 20 and 10% downsampled N0030. This parameter is no longer used in smCounter2, where the singletons are included if rpu $< 2$, excluded at random with probability $p = \text{rpu} - 2$ if $2 \leq \text{rpu} < 3$, and excluded if rpu $\geq 3$.

```
python sm_counter.py --outPrefix my_output_prefix --bamFile my_aligned_reads.bam --bedTarget
    my_target_region.bed --mtDepth mean_site_wise_UMI_depth --rpb mean_read_pairs_per_UMI --nCPU 64
     --minBQ 25 --minMQ 50 --hpLen 8 --mismatchThr 6.0 --mtDrop 0 or 1 --maxMT 0 --primerDist 2 --
```

```
threshold 0 --refGenome my_reference_genome.fa --bedTandemRepeats simpleRepeat.full.bed --
    bedRepeatMaskerSubset SR_LC_SL.full.bed --runPath /my/path --logFile my_output_prefix
```

The consensus reads were generated using `fgbio`'s `CallMolecularConsensusReads` (`min-reads` consistent with smCounter)

```
java -jar -Dsamjdk.use_async_io_write_samtools=true -Dsamjdk.use_async_io_read_samtools=true -
    Dsamjdk.compression_level=1 fgbio-0.1.4-SNAPSHOT.jar CallMolecularConsensusReads -i input.
    sorted.bam -o consensus.bam --min-reads=1 --min-input-base-quality=25 -1 40 -2 40 -D -S
    unsorted --tag=Mi
```

After necessary modifications to the headers, the consensus BAM file was filtered using `FilterConsensusReads`

```
java -jar fgbio-0.1.4-SNAPSHOT.jar FilterConsensusReads -i consensus.bam -o consensus.filtered.bam
    -r my_reference_genome.fa --min-reads=1 --max-base-error-rate=0.2 --min-base-quality=0
```

The MuTect command is

```
java -Xmx50g -jar muTect-1.1.4.jar --analysis_type MuTect --reference_sequence my_reference_genome.
    fa --intervals my_target_region.bed --input_file:consensus.filtered.bam --out
    mutect_extended_stats.out --coverage_file mutect_extended.wig.txt --vcf mutect_extended.vcf --
    enable_extended_output --fraction_contamination 0.003 --minimum_mutation_cell_fraction 0.004 --
    heavily_clipped_read_fraction 0.75 --min_qscore 20 --num_threads 20 --gap_events_threshold
    gapThr --downsample_to_coverage 200000
```

The VarDict command assuming minimum allele frequency of 0.5% is

```
VarDict -th 20 -F 0 -q 20 -G my_reference_genome.fasta -f 0.005 -N my_output_prefix -b consensus.
    filtered.bam -z -c 1 -S 2 -E 3 my_target_region.bed | teststrandbias.R | var2vcf_valid.pl -N
    my_output_prefix -E -f 0.005 > Variant_Call_Set.vcf
```

The HaplotypeCaller command is

```
java -jar GenomeAnalysisTK.jar -R my_reference_genome.fa -T HaplotypeCaller -stand_call_conf 30 -I
    consensus.filtered.bam -L my_target_region.bed -D dbSNP.vcf -o my_output.vcf
```

We varied MAGERI parameter `-defaultOverseq` from 4 to 16 to generate the ROC curve and used the default value for other parameters. The MAGERI command is

```
java -Xmx200g -jar mageri.jar --import-preset my_preset.xml -M4 --references my_reference_genome.fa
     --bed my_target_region.bed -R1 my_R1.fastq.gz -R2 my_R2.fastq.gz --project-name
    my_output_prefix -O /my/output/dir
```

### 4 Supplementary Figures

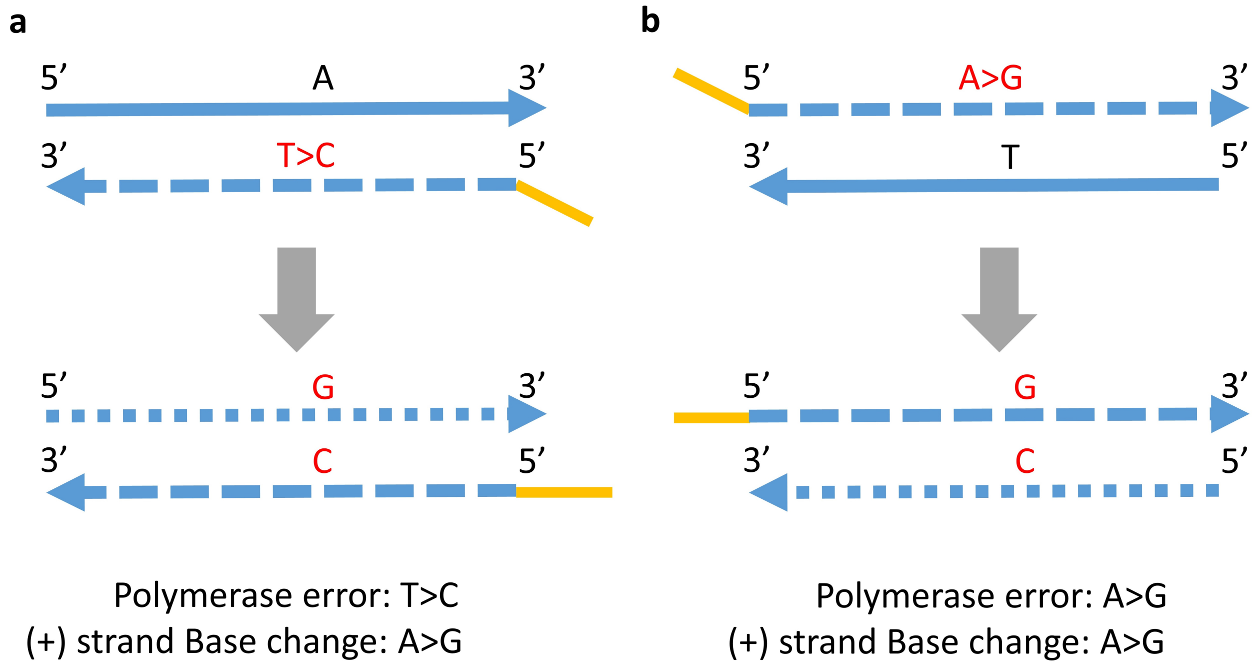


Figure S1: Counting DNA polymerase errors by strand. The same base change can be caused by two different polymerase errors, depending on the which DNA strand the primer binds to. **a** A polymerase incorporation error of T>C occurs when the primer binds to the forward strand of the original template (solid line). The final product is a double strand DNA with G and C on the forward (dotted line) and reverse (dashed line) strands , showing an A>G base change. **b** A polymerase incorporation error of A>G occurs when the primer binds to the reverse strand of the original template (solid line). The final product is a double strand DNA with G and C on the forward (dashed line) and reverse (dotted line) strands, also showing an A>G base change.

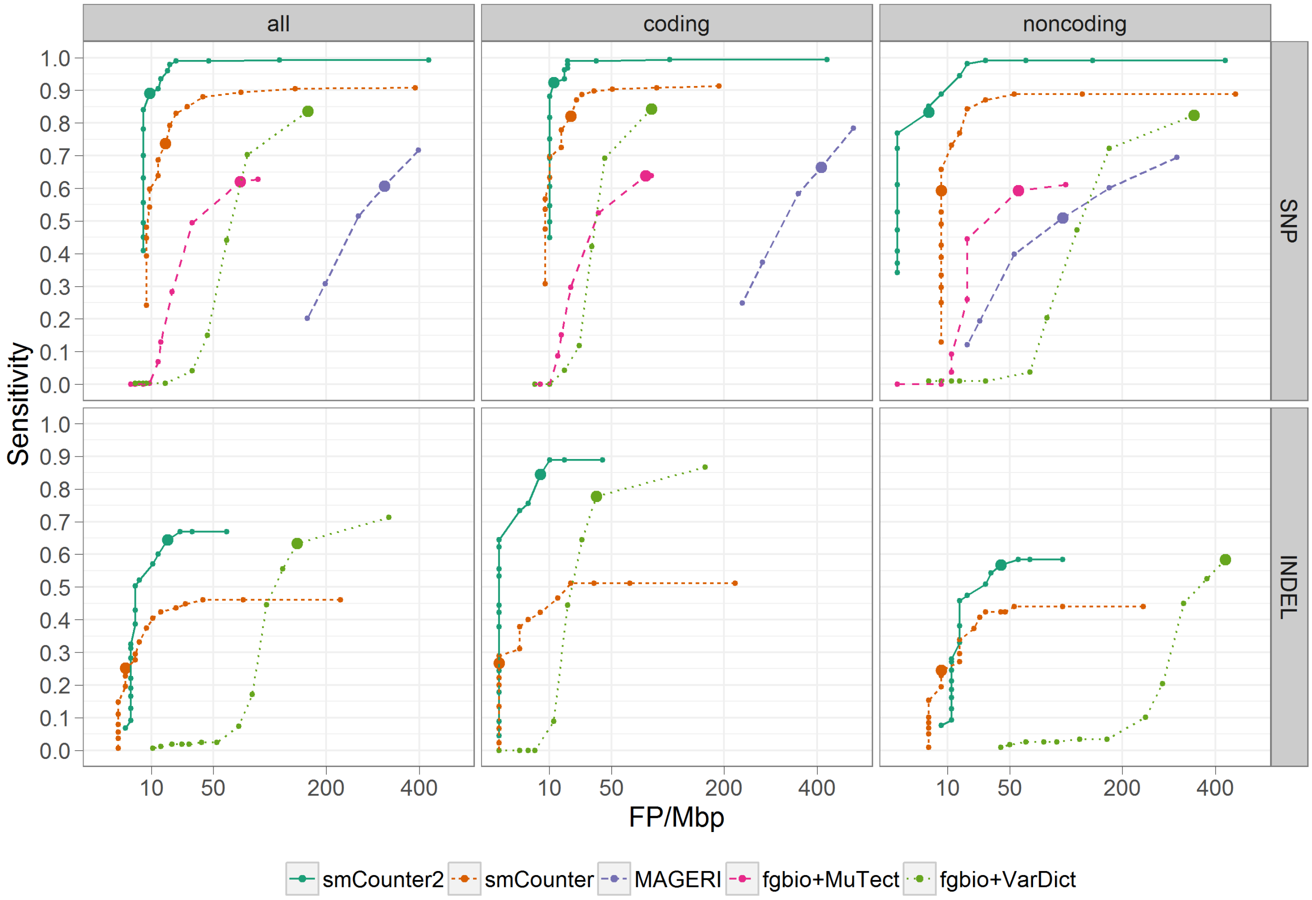


Figure S2: Benchmarking smCounter2, smCounter, fgbio+MuTect, fgbio+VarDict, and MAGERI on 0.5% variants in N13532. The performance is measured by false positives per megabase (x-axis) and sensitivity (y-axis), stratified by type of variant (SNV and indel) and genomic region (coding, non-coding, and all). The ROC curves are generated by varying the threshold for each method: Q-score for smCounter2, prediction index for smCounter, likelihood ratio for MuTect, minimum allele frequency for VarDict, and read per UMI for MAGERI while holding Q-score at 100. MuTect does not detect indels so is not included in the indel comparison. MAGERI did not call any indels correctly because of a bug that resulted in a one-base shift of indel positions.

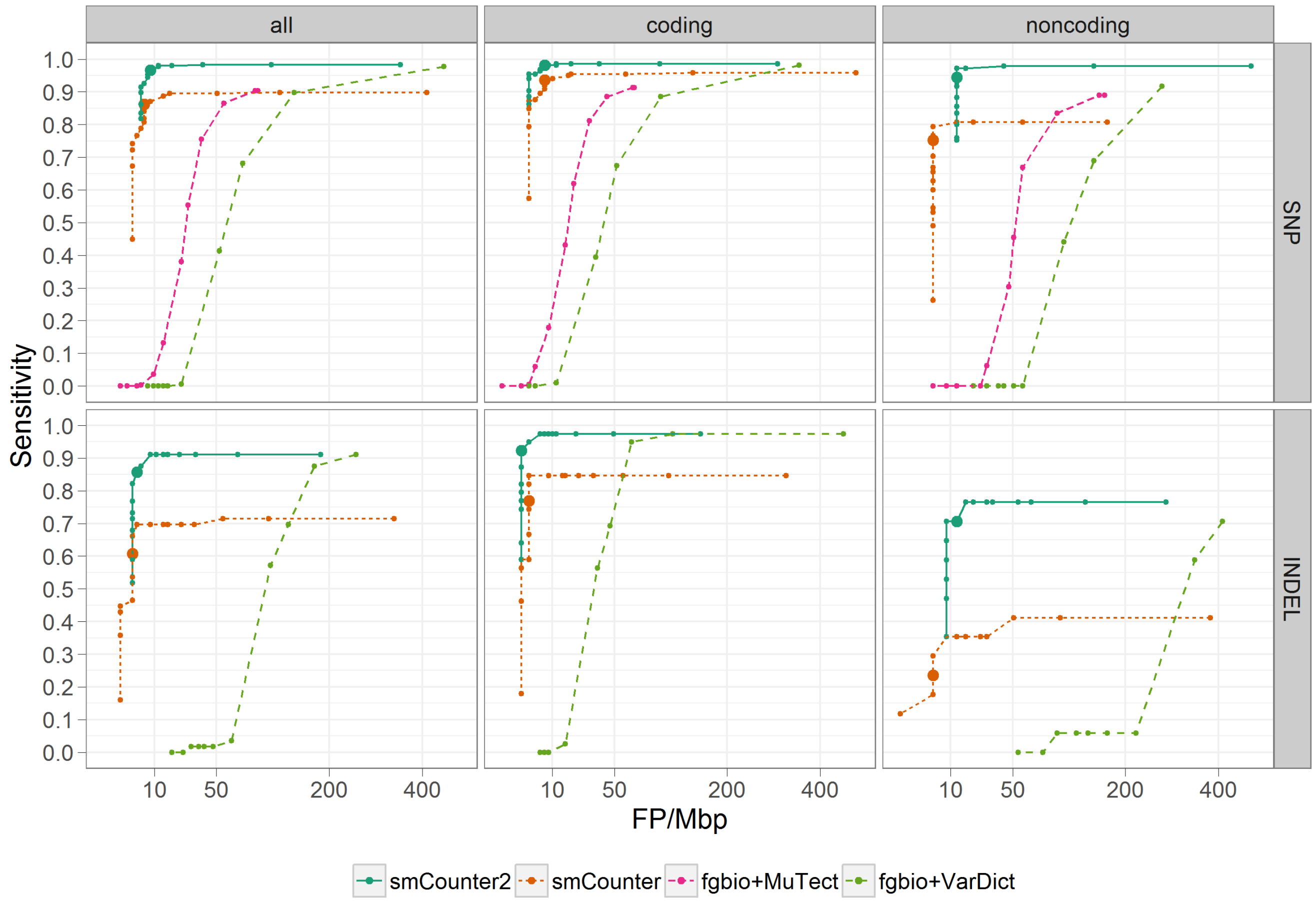


Figure S3: Benchmarking smCounter2, smCounter, fgbio+MuTect, and fgbio+VarDict on 1% variants in N0030. The performance is measured by false positives per megabase (x-axis) and sensitivity (y-axis), stratified by type of variant (SNV and indel) and genomic region (coding, non-coding, and all). The ROC curves are generated by varying the threshold for each method: Q-score for smCounter2, prediction index for smCounter, likelihood ratio for MuTect, and minimum allele frequency for VarDict. MuTect does not detect indels so is not included in the indel comparison.

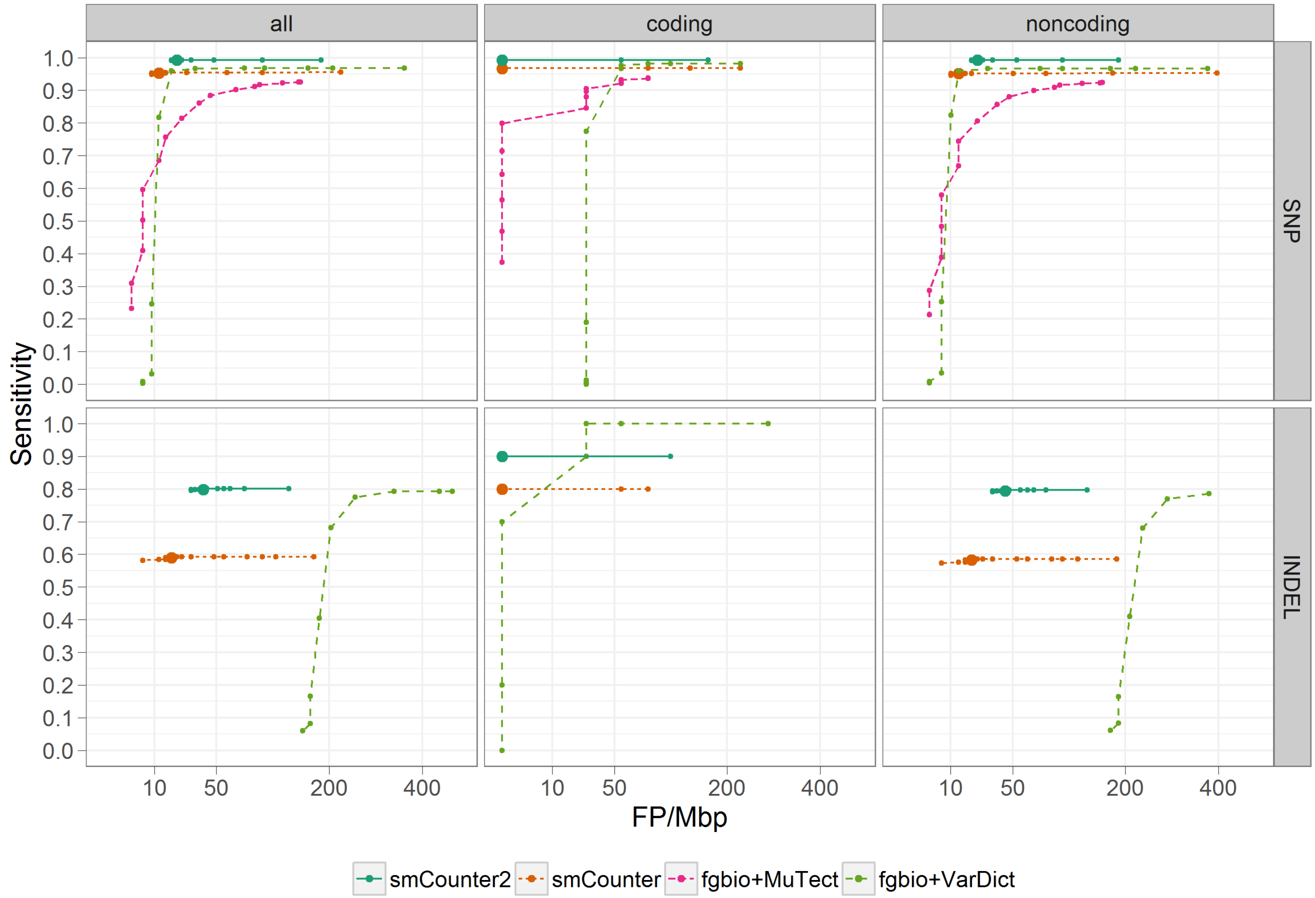


Figure S4: Benchmarking smCounter2, smCounter, fgbio+MuTect, and fgbio+VarDict on 5% variants in N0015. The performance is measured by false positives per megabase (x-axis) and sensitivity (y-axis), stratified by type of variant (SNV and indel) and genomic region (coding, non-coding, and all). The ROC curves are generated by varying the threshold for each method: Q-score for smCounter2, prediction index for smCounter, likelihood ratio for MuTect, and minimum allele frequency for VarDict. MuTect does not detect indels so is not included in the indel comparison.

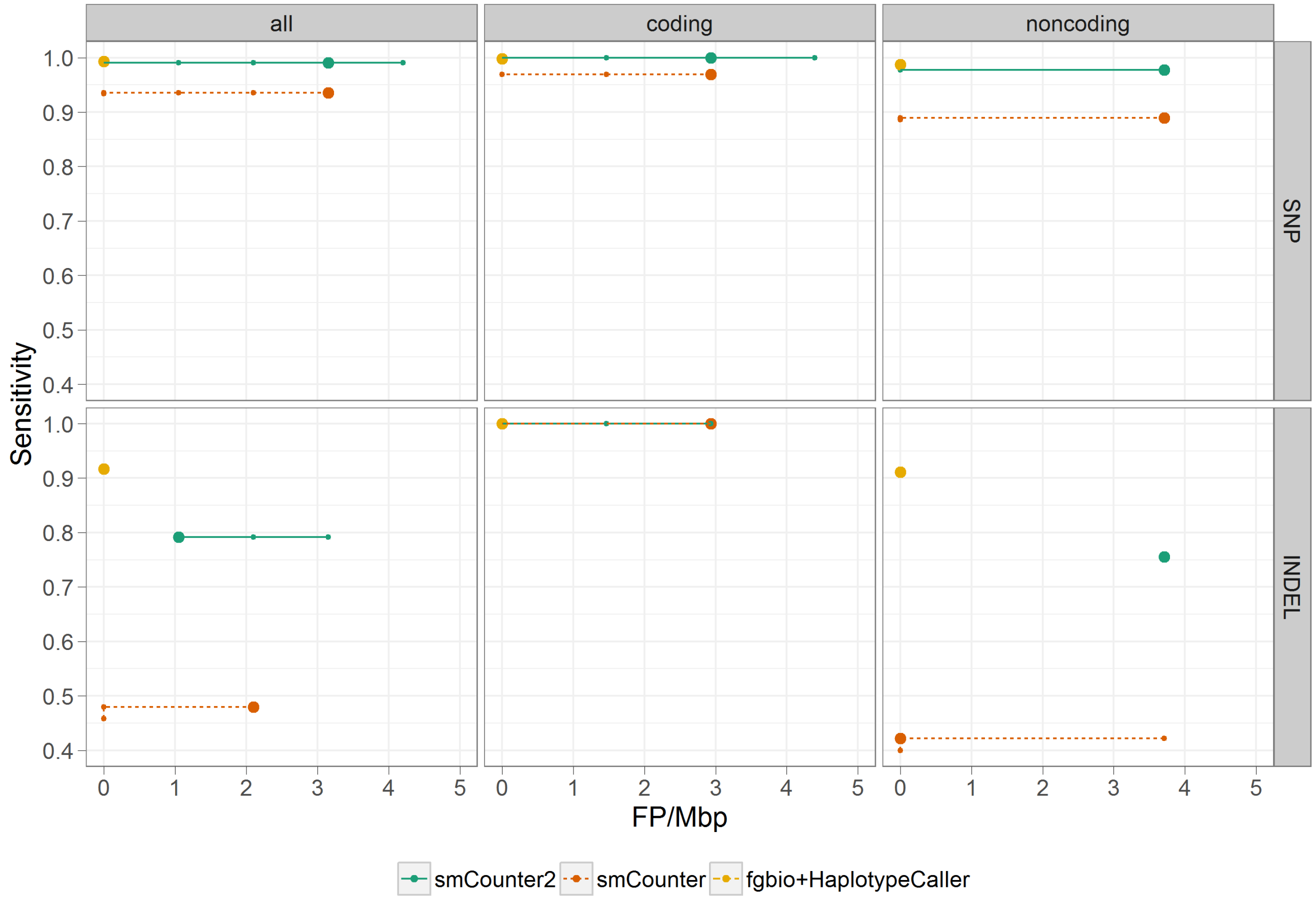


Figure S5: Benchmarking smCounter2, smCounter, fgbio+HaplotypeCaller on 50 and 100% variants in N11582. The performance is measured by false positives per megabase (x-axis) and sensitivity (y-axis), stratified by type of variant (SNV and indel) and genomic region (coding, non-coding, and all). The ROC curves are generated by varying the threshold for each method: Q-score for smCounter2, prediction index for smCounter, and minimum phred-scaled confidence threshold for HaplotypeCaller.

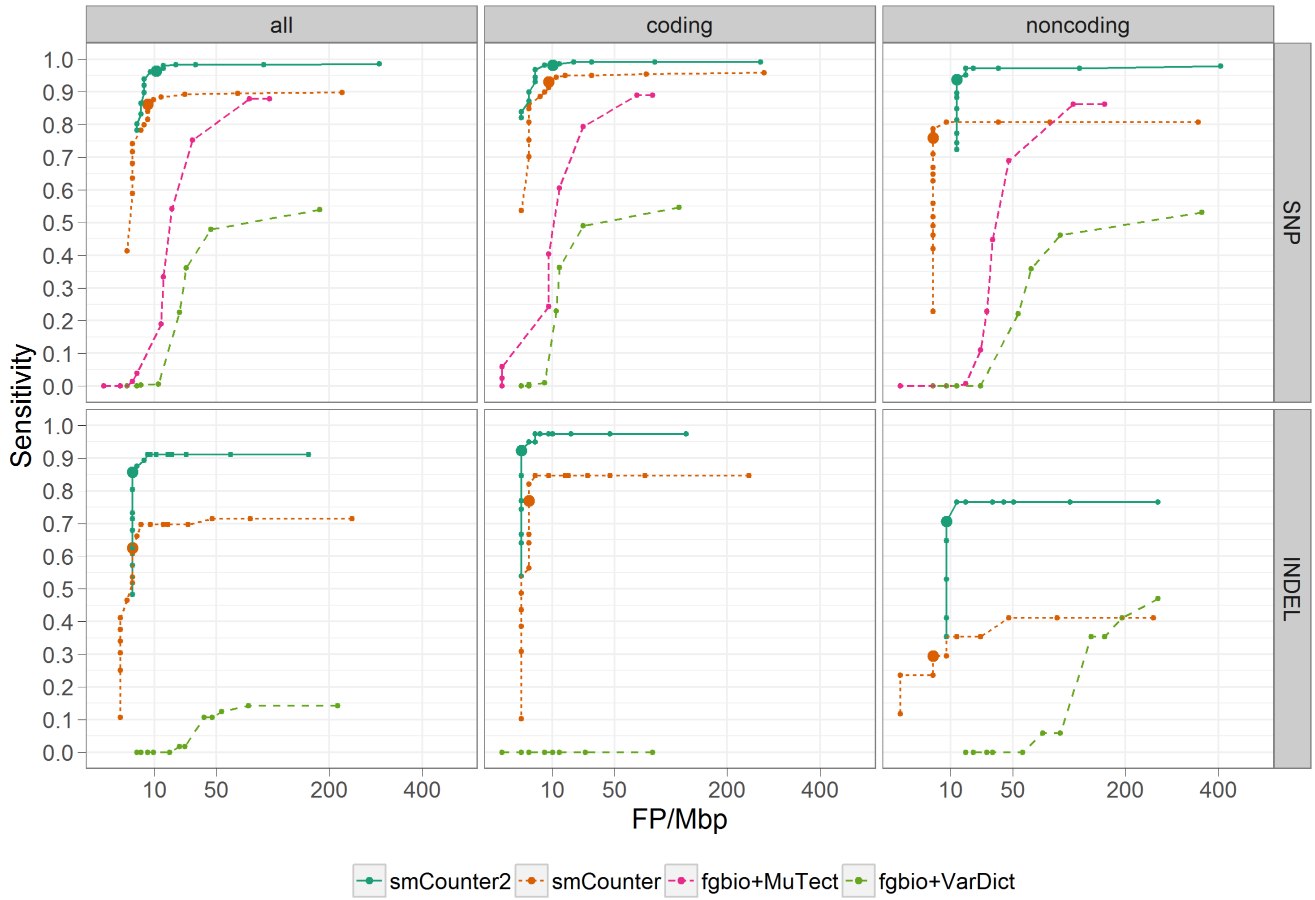


Figure S6: 80% downsampled N0030 (mean UMI depth: 1,742, rpu: 1.9) for benchmarking of smCounter2, smCounter, fgbio+MuTect, and fgbio+VarDict on 1% variants. The performance is measured by false positives per megabase (x-axis) and sensitivity (y-axis), stratified by type of variant (SNV and indel) and genomic region (coding, non-coding, and all). The ROC curves are generated by varying the threshold for each method: Q-score for smCounter2, prediction index for smCounter, likelihood ratio for MuTect, and minimum allele frequency for VarDict. MuTect does not detect indels so is not included in the indel comparison.

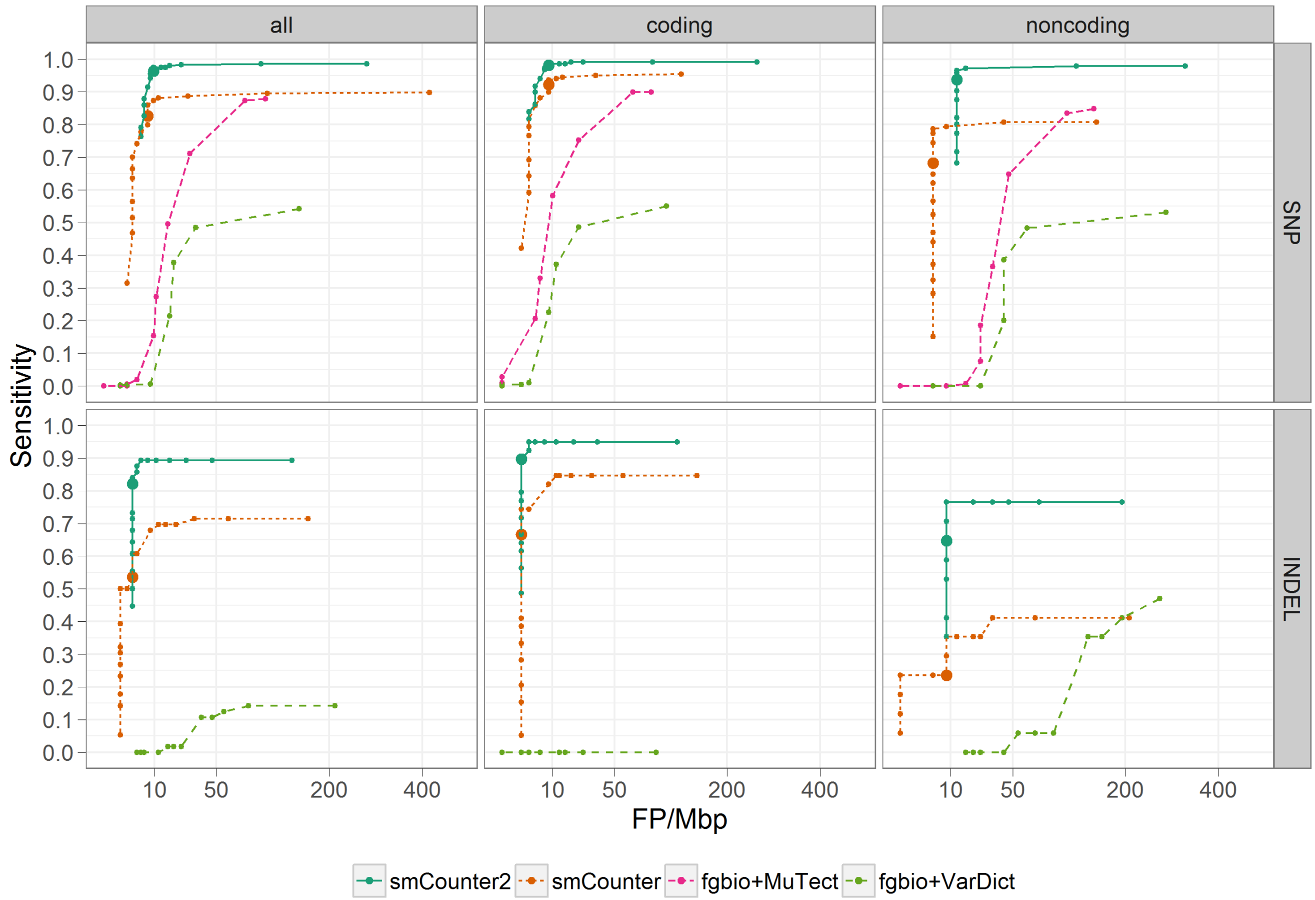


Figure S7: 60% downsampled N0030 (mean UMI depth: 2,362, rpu: 2.7) for benchmarking of smCounter2, smCounter, fgbio+MuTect, and fgbio+VarDict on 1% variants. The performance is measured by false positives per megabase (x-axis) and sensitivity (y-axis), stratified by type of variant (SNV and indel) and genomic region (coding, non-coding, and all). The ROC curves are generated by varying the threshold for each method: Q-score for smCounter2, prediction index for smCounter, likelihood ratio for MuTect, and minimum allele frequency for VarDict. MuTect does not detect indels so is not included in the indel comparison.

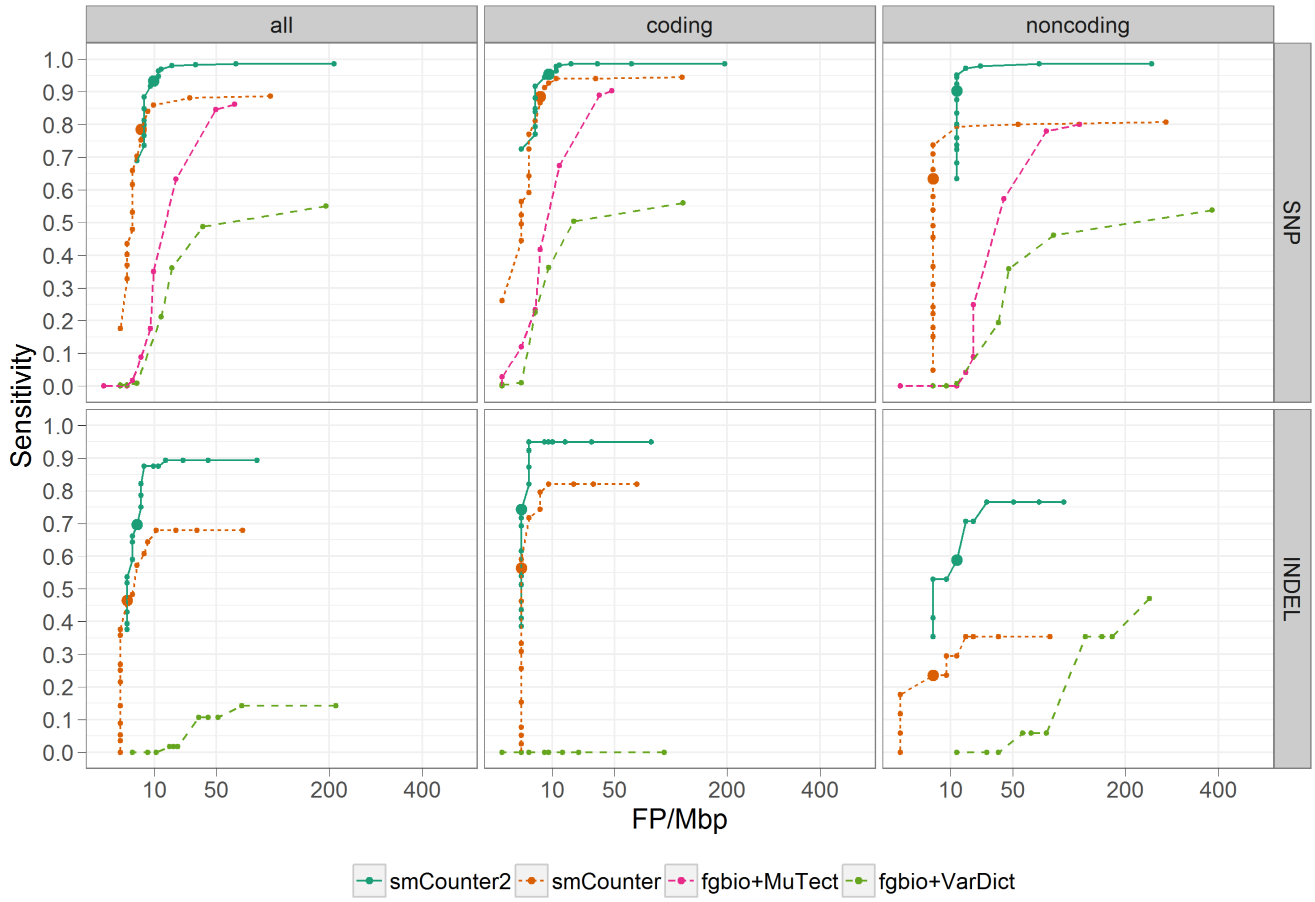


Figure S8: 40% downsampled N0030 (mean UMI depth: 2,943, rpu: 2.7) for benchmarking of smCounter2, smCounter, fgbio+MuTect, and fgbio+VarDict on 1% variants. The performance is measured by false positives per megabase (x-axis) and sensitivity (y-axis), stratified by type of variant (SNV and indel) and genomic region (coding, non-coding, and all). The ROC curves are generated by varying the threshold for each method: Q-score for smCounter2, prediction index for smCounter, likelihood ratio for MuTect, and minimum allele frequency for VarDict. MuTect does not detect indels so is not included in the indel comparison.

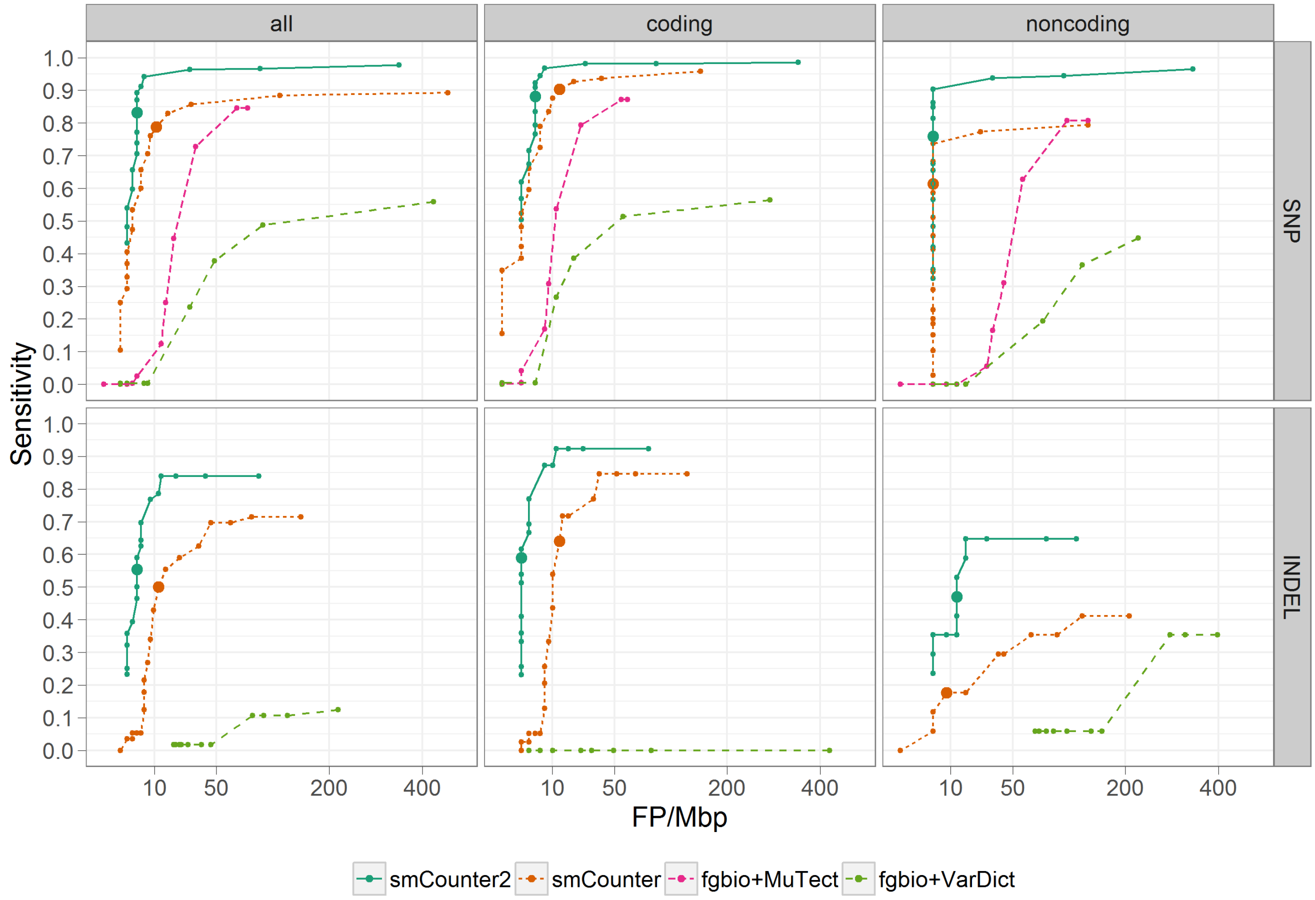


Figure S9: 20% downsampled N0030 (mean UMI depth: 3,252, rpu: 5.8) for benchmarking of smCounter2, smCounter, fgbio+MuTect, and fgbio+VarDict on 1% variants. The performance is measured by false positives per megabase (x-axis) and sensitivity (y-axis), stratified by type of variant (SNV and indel) and genomic region (coding, non-coding, and all). The ROC curves are generated by varying the threshold for each method: Q-score for smCounter2, prediction index for smCounter, likelihood ratio for MuTect, and minimum allele frequency for VarDict. MuTect does not detect indels so is not included in the indel comparison.

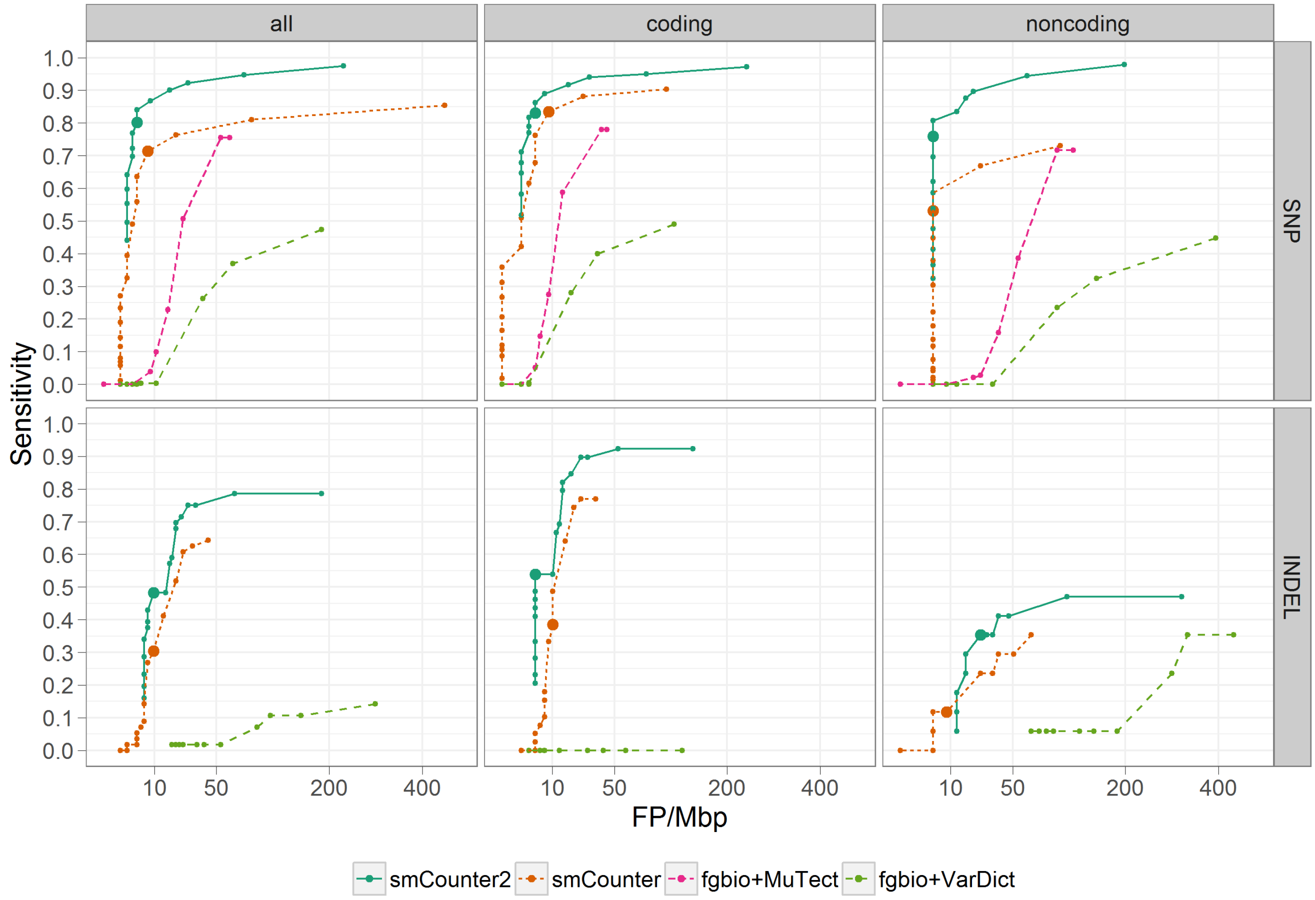


Figure S10: 10% downsampled N0030 (mean UMI depth: 3,454, rpu: 7.2) for benchmarking of smCounter2, smCounter, fgbio+MuTect, and fgbio+VarDict on 1% variants. The performance is measured by false positives per megabase (x-axis) and sensitivity (y-axis), stratified by type of variant (SNV and indel) and genomic region (coding, non-coding, and all). The ROC curves are generated by varying the threshold for each method: Q-score for smCounter2, prediction index for smCounter, likelihood ratio for MuTect, and minimum allele frequency for VarDict. MuTect does not detect indels so is not included in the indel comparison.
